# Attenuation of polysialic acid biosynthesis in cells by the small molecule inhibitor 8-keto-sialic acid

**DOI:** 10.1101/2022.09.02.506362

**Authors:** Carmanah Hunter, Zhizeng Gao, Hong-Ming Chen, Nicole Thompson, Warren Wakarchuk, Mark Nitz, Stephen G. Withers, Lisa M. Willis

**Affiliations:** Department of Biological Sciences, University of Alberta, Edmonton, T6G 2R3, Canada; Department of Chemistry, University of British Columbia, Vancouver, V6T 1Z1, Canada; Department of Chemistry, University of Toronto, Toronto, M5S 3H6, Canada

**Keywords:** polysialic acid, polysialyltransferase, metabolic oligosaccharide engineering, Neu5Ac analog, sialyltransferase inhibitors, glycoengineering

## Abstract

Sialic acids are key mediators of cell function, particularly with regards to cellular interactions with the surrounding environment. Reagents that modulate the display of specific sialyl glycoforms at the cell surface would be useful biochemical tools and potentially allow for therapeutic intervention in numerous challenging chronic diseases. While multiple strategies are being explored for the control of cell surface sialosides, none that shows high selectivity between sialyltransferases or that targets a specific sialyl glycoform has yet to emerge. Here we describe a strategy to block the formation of α2,8-linked sialic acid chains (oligo- and polysialic acid) through the use of 8-keto-sialic acid as a chain-terminating metabolic inhibitor that, if incorporated, can not be elongated. 8-Keto-sialic acid is non-toxic at effective concentrations and serves to block polysialic acid synthesis in cancer cell lines and primary immune cells, with minimal effects on other sialyl glycoforms.

## Introduction

*N*-Acetylneuraminic acid (Neu5Ac), the most abundant sialic acid, often terminates the glycan structures of the glycocalyx surrounding all human cells.[1] Cell signaling interactions and pathogen recognition are commonly mediated through binding events with glycoproteins and glycolipids that display Neu5Ac.[2] Some of the best studied examples of Neu5Ac-mediated interactions include the recognition of cell surfaces by viral hemagglutinin in influenza infections and the rolling of leukocytes at the site of inflammation.[3, 4] Sialyl glycoforms are diverse, with sialyltransferase enzymes catalyzing the attachment of Neu5Ac to underlying *N*- and *O*-linked glycans through α2,3- or α2,6-linkages to galactose and *N*-acetylgalactosamine residues, and through an α2,8-linkage to Neu5Ac.[1] Aberrant sialylation is a hallmark of numerous chronic diseases, including autoimmune diseases and cancers and knock out mouse studies of specific sialyltransferases have demonstrated the potential therapeutic effects of modulating sialyltransferase activity.[5–8] Thus, methods for modulation of specific sialyl glycoforms would be a useful pharmacological tools and may provide strategies for therapeutic intervention in numerous challenging chronic diseases.

Multiple strategies are being explored for the control of cell surface sialosides. Direct inhibition of the sialyltransferase enzymes has received considerable attention, although few studies have progressed beyond *in vitro* characterization.[9] Recently, inhibitor scaffolds based on CMP derivatives functionalized with amino-phenyl-hydroxymethyl phosphonic acids have enabled the development of cell-permeable pan-selective sialyltransferase inhibitors.[10, 11] Similarly, methylated CMP derivatives have been found to have some selectivity in sialyltransferase inhibition both *in vitro* and in cell culture.[12] Metabolic strategies that exploit the endogenous monosaccharide salvage pathways to reduce global sialylation levels have also been effective. Treatment of both cells and mice with 3F_ax_-Neu5Ac results in production of CMP-3F_ax_-Neu5Ac, which is a competitive inhibitor of sialyltransferases and also causes negative feedback inhibition for the synthesis of CMP-Neu5Ac.[13, 14] Alternatively, reduction of the cellular concentration of CMP-Neu5Ac has been achieved with 3-OMe ManNAc, which inhibits mannosamine kinase, thereby blocking sialic acid biosynthesis.[15]

While these approaches show promise, a strategy that shows high selectivity between sialyltransferases or targets a specific sialyl glycoform has yet to emerge. In particular, we were interested in strategies to selectively block biosynthesis of the important glycan polysialic acid (polySia). This linear polymer of α2,8-linked Neu5Ac can be up to 400 residues long and confers profound consequences on the proteins and cells on which it is found. In healthy adults, its expression is limited to a small subset of proteins found in the nervous, immune, and reproductive systems but it is also strongly overexpressed in many cancers.[7] Like many other sialosides, polySia acts as an immune checkpoint in that it attenuates the immune response. Depending on the cell in which it is expressed, polySia can decrease phagocytosis, cell proliferation, and the release of proinflammatory cytokines.[16–18] In the case of cancer, polySia is widely expressed in many cancer cells and the link between polySia and metastasis is well established.[6, 19–21] *In vitro* models have demonstrated that polySia promotes cell migration [22–25] and its importance for cell migration and metastasis has been established in *in vivo* models.[20, 24, 26, 27] Additionally, high expression levels of polySia in clinical samples correlate with significantly increased metastasis and poor prognoses.[6]

Selective tools to block the formation of polySia in both primary immune and cancer cells would accelerate experiments to reveal how this glycan functions in health and disease and possibly lead to new therapeutic approaches based on lowering polySia levels. A logical strategy might be to synthesize 8-deoxy analogues of sialic acid as chain-terminating metabolic inhibitors that could be incorporated but not elongated. Unfortunately synthetic routes to such analogues are lengthy [28]. An attractive alternative is the sterically conservative 8-keto substituent (Figure 1), which could still engage in hydrogen bonding interactions yet not act as an acceptor. Importantly, synthesis of this analogue is enabled by a simple palladium catalyst that facilitates oxidation exclusively at the 8-position.[29] We hypothesized that metabolic incorporation of 8-keto-Neu5Ac could selectively reduce the formation of polySia while maintaining the synthesis of α2,3- and α2,6-linkages. Here, we demonstrate that 8-keto-Neu5Ac is a metabolic glycoengineering tool that is non-toxic at working concentrations and reduces polySia synthesis in both cancer cell lines and primary immune cells with minimal effects on other sialiosides.

**Figure 1.**
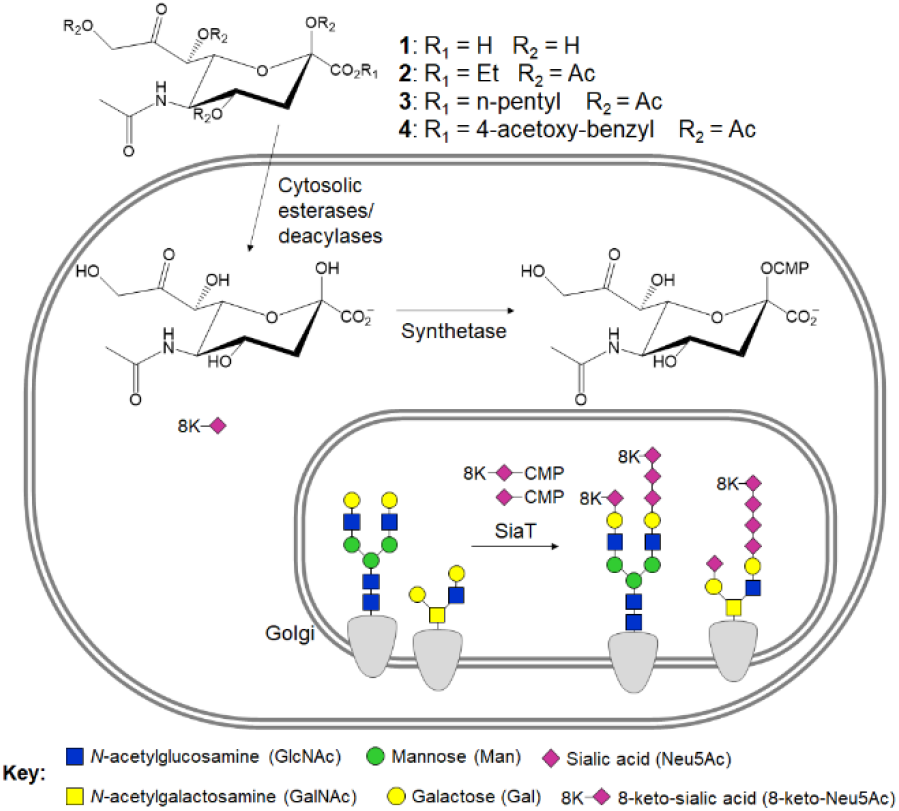
8-Keto-Neu5Ac as an inhibitor of α2,8-linked sialyl glycoforms. Esterified and per-*O*-acetylated 8-keto-Neu5Ac is taken up by cells and converted to the corresponding nucleotide sugars, which are incorporated into sialyl glycoforms. The 8-keto moiety is not a substrate for further extension of α2,8-linkages.

## Results

### 8-Keto-Neu5Ac derivatives block polysialylation in cancer cell lines

The 8 position of Neu5Ac benzyl ester was selectively oxidized using the previously reported Pd catalyst in a single step which, following hydrogenolysis, afforded 8-keto-Neu5Ac (**1**) (See SI for details).[29] This was then converted into esterified versions to improve cell permeability. Based upon our previous exploration of ester pro-drug versions of modified sialic acids [30], we prepared the ethyl, n-pentyl, and 4-acetoxybenzyl esters at C-1, each in their otherwise per-*O*-acetylated forms. The effect of per-*O*-acetylated 8-keto-Neu5Ac esters on polysialic acid expression was assessed in both MCF-7 and AtT-20 cancer cell lines. MCF-7 is an adherent breast cancer cell line that makes a small amount of intracellular (Golgi-localized) polySia [31] and AtT-20, which grow as suspended clusters, is a pituitary cancer cell line that makes large amounts of cell surface polySia [32]. After three days of incubation with **2-4**, polySia was examined by α-polySia immunoblot of cell lysates and quantified by densitometry (Figure 2A-D). All three esters caused a dose-dependent decrease in polySia in MCF-7 cells, with **3** having the strongest effect. **3** was also the only derivative to substantially affect polySia in AtT-20 cells. In both cell lines, 100 μM **3** was sufficient to reduce polySia by at least 50%. No major trend in reduction of polysialic acid length, as determined by a shift in gel mobility, was observed with increasing concentrations of the 8-keto-Neu5Ac derivatives suggesting preferential blockade of early steps in the polymerization, as has been seen with other polysialyltransferase inhibitors. [33]

**Figure 2.**
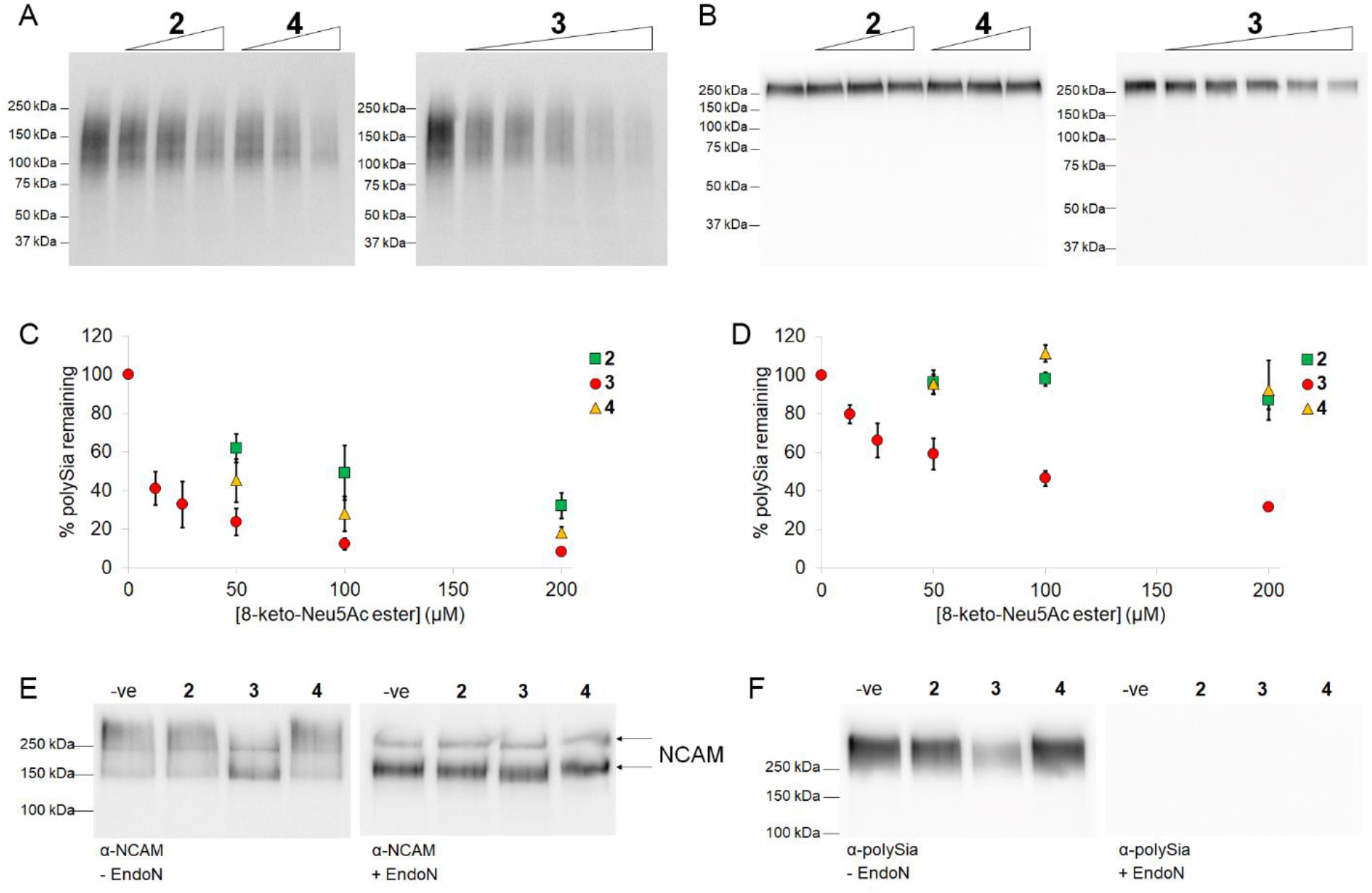
Inhibition of polysialic acid biosynthesis by 8-keto-Neu5Ac derivatives. α-PolySia immunoblots of (A) MCF7 and (B) AtT-20 cells treated with 8-keto-Neu5Ac esters for three days. Equivalent amounts of protein were loaded in each well. Quantification of polySia in (C) MCF7 and (D) AtT-20 using densitometry of blots in (A) and (B) respectively. Immunoblots showing NCAM (E) and polySia (F) from AtT-20 cell lysates after treating cells with 100 μM 2, 3 and 4 for three days. Cells were treated +/− the polySia-specfic endosialidase (EndoN) prior to loading equivalent amounts of protein on the gel.

To ensure that the observed reduction in global polysialylation upon treatment with **2-4** reflected only loss of the glycoform and not loss of protein polysialylation substrates, the effect of 8-keto-Neu5Ac esters on a specific protein known to be polysialylated was examined. NCAM is the major polysialylated protein in AtT-20 cells and polysialylation of NCAM causes a shift in apparent molecular weight of the protein. Cells treated with **2**-**4** (100 μM) have the same amount of NCAM as the untreated control (Figure 2E, right panel), indicating that the reduction in polysialylation is not a result of decrease in expression of the underlying protein. Incubation of AtT-20 cells with 100 μM **2** and **4** resulted in no change to the migration of NCAM (Figure 2E, left panel), consistent with the observation that neither of these esters decreases polySia in AtT-20 cells at this concentration (Figure 2B,F). In contrast, 100 μM **3** causes a major shift in the migration of NCAM which is similar to the shift observed upon enzymatic removal of polySia using the polysialic acid-specific endoglycosidase EndoN. The similar banding pattern seen between treatment with **3** and EndoN suggests the observed loss of polySia signal is not due to a loss of reactivity with the α-polySia antibody against the 8-keto-Neu5Ac modified polySia, as can occur with other modifications, such as *N*-butanoylmannosamine.[34, 35]

To confirm that the 8-keto-Neu5Ac derivatives were not toxic, we measured cell viability in both cell lines. In MCF-7 cells, **4** reduced their viability, likely due to the generation of a toxic quinone methide upon deacetylation of the 4-acetoxybenzyl ester, while **2** and **3** were approximately 10 times less damaging (Table 1, S1). In contrast, none of the compounds displayed any toxicity towards AtT-20 cells, even at concentrations up to 4 mM, the highest concentration tested (Table 1, S1).

**Table 1.** Toxicity of 8-keto-Neu5Ac derivatives. IC50 values for 8-keto-Neu5Ac esters were measured using a WST-1 assay after 24 h incubation with human breast (MCF7) and rat pituitary (AtT-20) cancer cell lines.

|  | **2** | **3** | **4** |
| --- | --- | --- | --- |
| MCF7 | ~3 mM | ~3 mM | 0.3 mM |
| AtT-20 | >4 mM | >4 mM | >4 mM |

### 8-Keto-Neu5Ac disrupts polySia synthesis through polySia chain termination

Given our goal of making a glycoengineering tool that selectively perturbs the synthesis of α2→8 linked sialic acids, it was necessary to determine whether treatment with 8-keto-Neu5Ac interrupted the synthesis of other sialyl glycoforms. Our hypothesis when designing this metabolic glycoengineering tool was that 8-keto-Neu5Ac would not act as an inhibitor of sialyltransferases or CMP-Neu5Ac synthases, but rather would be incorporated into sialyl glycoforms and act as a polySia chain terminator preventing elongation of polySia chains by α2→8 sialyltransferases. As such, we expected 8-keto-Neu5Ac (**1**) to be a substrate for CMP-Neu5Ac synthetases and sialyltransferases.

To test the suitability of **1** as a substrate for sialyltransferases *in vitro*, it was first converted to CMP-8-keto-Neu5Ac using the previously reported bacterial CMP-Neu5Ac synthetase [36] (Figure S2). CMP-8-keto-Neu5Ac was evaluated as a substrate for three recombinant mammalian sialyltransferases, ST3GalI, ST6GalI, and ST8SiaIV using fluorescently tagged synthetic T antigen (Gal-β1,3-GalNAc), LacNAc (Gal-β1,4-GlcNAc), and disialyl-Lac (Neu5Ac-α2,8-Neu5Ac-α2,3-Gal-β1,4-Glc) acceptors respectively (Figure S3). We were surprised to observe that although all three enzymes transferred CMP-8-keto-Neu5Ac to their respective substrates, they did so with substantially lower efficiency than they transferred the natural donor CMP-Neu5Ac (Figure 3A-C). To see if CMP-8-keto-Neu5Ac was acting as an inhibitor, we performed mixed donor sugar assays. For ST3GalI, although addition of 8-keto-Neu5Ac had no effect on product formation, addition of CMP-8-keto-Neu5Ac reduced the amount of product formed by ~60%, consistent with recognition and slow turnover of the modified substrate. While this reduction is significant, the concentrations of CMP-8-keto-Neu5Ac (1.5 mM) used are likely higher than would be generated *in cellulo* at working concentrations (200 μM). For ST6GalI and ST8SiaIV, addition of an equivalent concentration of CMP-8-keto-Neu5Ac or its precursor 8-keto-Neu5Ac had no effect on the ability of the enzyme to transfer Neu5Ac to its substrate, suggesting that CMP-8-keto-Neu5Ac is poorly recognized by these enzymes but not inhibiting them. Overall, the capacity for these enzymes to use CMP-8-keto-Neu5Ac as an (albeit poor) substrate, and the absence of inhibition of two of these enzymes in the mixed donor sugar assays supports 8-keto-Neu5Ac as a polySia chain terminator rather than an inhibitor.

**Figure 3.**
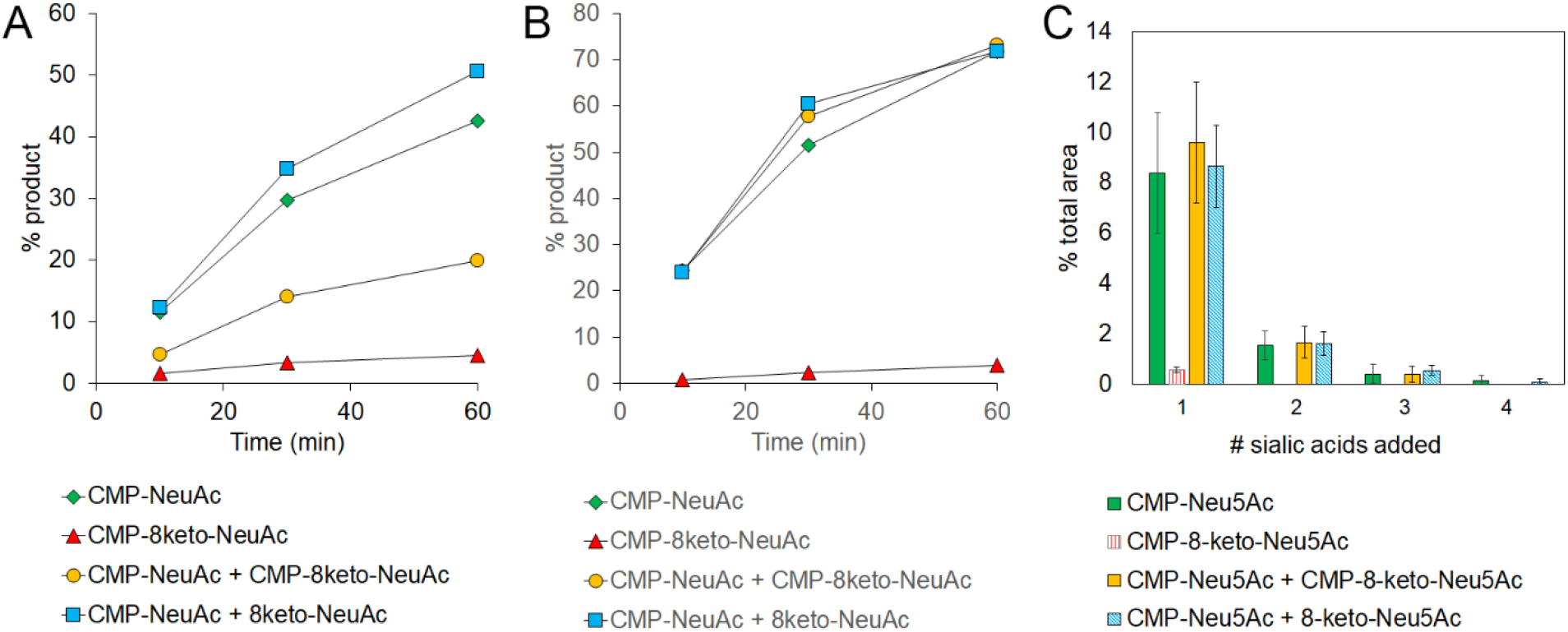
Sialyltransferases use CMP-8-keto-Neu5Ac as a substrate with less efficiency than CMP-Neu5Ac. Activity of ST3Gall (A), ST6Gall (B), and ST8SialV (C) with fluorescent T-antigen, LacNAc, and disialyl-Lac acceptors, respectively. 1.5 mM donor sugar was used for A and B, 2 mM donor sugar was used for C.

Another potential mechanism for blocking of sialic acid synthesis *in cellulo* is through negative feedback inhibition of the synthesis of CMP-Neu5Ac, resulting in a loss of cell surface sialic acid. This mechanism of action is shared by previous global inhibitors of sialylation, 3-F_ax_-Neu5Ac and CMP [13, 37]. Metabolic incorporation of **3** and of the corresponding 3-F_ax_-Neu5Ac pentyl ester in MCF7 cells both resulted in a reduction in cellular polySia (Fig 5A). If 8-keto-Neu5Ac were similarly inhibiting polysialylation through dysregulation of CMP-Neu5Ac synthesis, we would expect that all cell surface Neu5Ac would decrease upon metabolic incorporation of **3**. We examined whether metabolic incorporation of **3** disrupted cell surface Neu5Ac by labeling the surface of MCF7 cells with lectins followed by analysis with flow cytometry. To determine whether α2,3-linked Neu5Ac was minimally perturbed in cells, we investigated the effect of compound **3** on the ability of MAL-II, an α2,3-linked Neu5Ac-specific lectin, to label cells (Figure 4B). This analysis was particularly important given our observation that high concentrations of CMP-8-keto-Neu5Ac partially inhibited recombinant ST3GalI *in vitro*. MCF-7 cells treated with 200 μM compound **3** for three days were collected, labeled with biotinylated MAL-II and streptavidin-PE, and analyzed by flow cytometry. As a control, we treated cells, separately, with 200 μM of the corresponding peracetylated 3F_ax_-Neu5Ac pentyl ester or with a non-specific neuraminidase NedA to remove most sialic acids. No significant differences in MAL-II staining were observed in cells treated with 8-keto-Neu5Ac, but reductions in staining were observed for both 3F_ax_-Neu5Ac and neuraminidase-treated cells, confirming that at working concentrations compound **3** does not inhibit cell surface α2,3-sialylation. Probing the MCF-7 cell surface with SNA, a lectin specific for α2,6-linked sialic acids, was unsuccessful (data not shown) which is consistent with other reports suggesting that MCF7 cells do not display many α2,6-linked sialic acids on their cell surface [38]. However, staining with PNA, a lectin that recognizes galactose, complemented the results of the MAL-II staining (Figure 4C). While **3** did not have a significant effect on PNA staining, cells treated with 3F_ax_-Neu5Ac or neuraminidase stained more heavily, indicating more exposed galactose, thus fewer cell surface sialic acids. These experiments provide further evidence that **3** is not primarily interrupting cellular sialylation pathways through global disruption of CMP-Neu5Ac synthesis or global sialyltransferase inhibition, but rather that 8-keto-Neu5Ac is a metabolic glycoengineering tool that preferentially disrupts the synthesis of α2,8-linked sialic acids.

**Figure 4.**
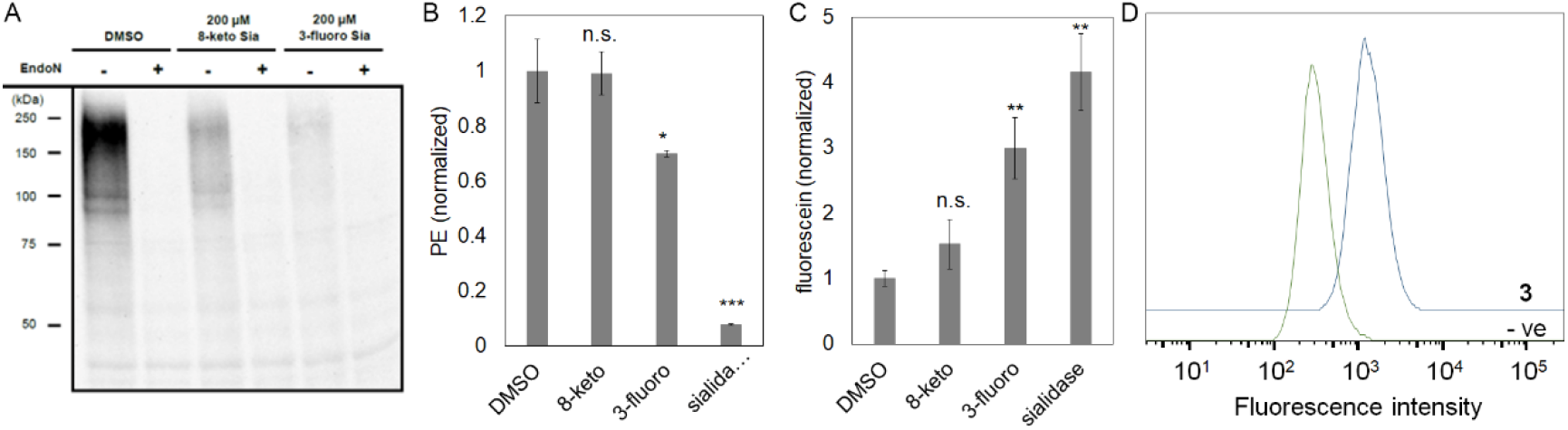
8-keto-Neu5Ac does not inhibit cell surface sialylation. MCF-7 cells were treated with 200 μM of **3** (8-keto-Neu5Ac) or 3-fluoro-Neu5Ac (3-fluoro) for three days and then analyzed for presence of polySia (A - α-polySia immu noblot), α-2,3-linked Neu5Ac (B - flow cytometry of cells labeled with biotinylated MAL-II, followed by streptavidin-PE), galactose (C - flow cytometry of cells labeled with fluorescein-conjugated PNA), and cell surface ketones (D - flow cytometry of cells treated with biotin-hydrazide followed by avidin-FITC). Both **3** (8-keto-NeuAc) and 3-fluoro-Neu5Ac contained a pentyl ester at C1 and were otherwise peracetylated.

Finally, we demonstrated that the 8-keto-Neu5Ac derivative **3** is metabolically processed for cell surface display via cellular sialylation pathways. MCF-7 cells were incubated for three days with 100 μM of **3** and then reacted with biotin-hydrazide to label the cell surface ketones. After treating cells with FITC-avidin, flow cytometry analysis revealed an increase in fluorescence for cells treated with **3** (Figure 4D). This data is consistent with our observations *in vitro* that CMP-8-keto-Neu5Ac was a substrate for recombinant sialyltransferases, and demonstrates that 8-keto-Neu5Ac is processed by both CMP-Neu5Ac synthases and sialyltransferases for cell surface display. Taken together, the evidence supports the model that 8-keto-Neu5Ac is a metabolic tool that functions as a polySia chain terminator preferential for the reduction of polySia over other sialoglycoforms.

### 8-Keto-Neu5Ac reversibly reduces polySia levels in primary T cells

While the ability to manipulate polySia in cultured cell lines is advantageous, we would ideally like to be able to modify polySia in primary cells, which are directly relevant to human biology but are notoriously difficult to manipulate genetically. To determine whether compound **3** was effective in primary cells, its effect on polySia was examined in activated human CD3^+^ T cells. T cells play a central role in cell-mediated immunity and while they contain polySia, the role of polySia in T cell activation and effector function has not yet been elucidated. Within 24 h, activated CD3^+^ T cells treated with 200 μM **3** showed a substantial decrease in polySia compared to the DMSO-treated control cells (Figure 5, S4). This decrease persisted until day three, when the cells were supplied with fresh media that did not contain 8-keto-Neu5Ac. By day five, polySia had recovered to the level of untreated cells. A similar restoration of polySia after consumption of 8-keto-Neu5Ac and polySia turnover was observed in MCF7 and Jurkat cells (Figure S5,S6), confirming that this reduction in polySia levels is dependent on the presence of 8-keto-Neu5Ac and that this probe does not permanently alter polySia metabolism.

**Figure 5.**
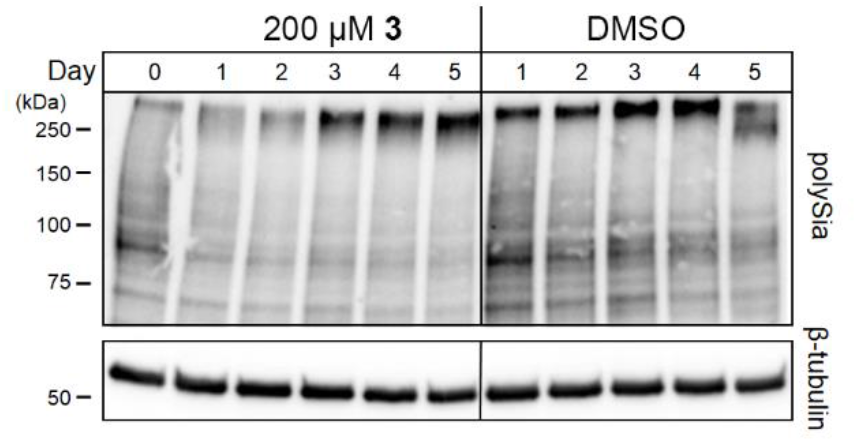
8-Keto-Neu5Ac reversibly reduces polySia levels in primary T cells. Activated human CD3* T cells were incubated for 3 days +/− 200 μM compound **3** (days 0-3), after which the cells were diluted to 5×10^5^/mL and incubated a further 2 days in media lacking 8-keto-Neu5Ac (days 4 and 5). Each day cells were collected, washed, and lysed. Equivalent amounts of protein were loaded on the gel and the blot was probed with α-polySia mAb and (β-tubulin as a loading control. Representative immunoblot shown.

## Discussion

We have developed an inhibitor of sialylation that is selective for sialyl glycoforms containing α2,8-linkages rather than the global reduction in sialylation observed with all other reported inhibitors. The type of ester protecting group makes a large difference in the reduction of polySia, suggesting that the limiting step is cytosolic deprotection of the sugar and that improvement in the cytosolic availability of the free sugar may then increase the potency of inhibition, as seen previously in a parallel system [30]. While modification of glycosyl acceptors, generally by methylation or deoxygenation, has been shown to be an effective strategy to inhibit glycosyltransferases in vitro, few such studies have been performed *in vivo* and to our knowledge, in no case has this involved a ketone substitution.[39, 40] With 8-keto-Neu5Ac, further extension at the 8 position is blocked. The hydrated ketone may be an acceptor but the resulting product would be an unstable hemiketal, likely undergoing elimination to the parent ketone with a half-life of seconds to minutes under neutral aqueous conditions based on the rates of ketone hydrate dehydration.[41]

Ideally, the 8-keto modification would be sufficiently similar to Neu5Ac to be a substrate for monosialylation by sialyltransferases and subsequent recognition by lectins and other sialic acid-binding proteins, minimally perturbing the biology of α2,3- or α2,6-linked sialyl glycoforms. However, we were surprised to see that 8-keto-Neu5Ac is a poor substrate for all three sialyltransferases tested. The only difference between CMP-Neu5Ac and CMP-8-keto-Neu5Ac, besides the loss of a hydrogen bond donor, is the change in geometry at position 8 from sp^3^ to sp^2^ hybridization and the possible presence of some of the ketone hydrate form. These results highlight the importance of the glycerol side chain to sialyltransferase binding. However, the reduced activity of sialyltransferases towards 8-keto-Neu5Ac may actually be of benefit to our overall goal of producing a minimally perturbing inhibitor of α2,8-linked Neu5Ac; as monosialyltransferases prefer the natural donor sugar, much of the α2,3- and α2,6-linked sialyl glycoforms are likely to be Neu5Ac. Regardless of whether sialyl glycoforms contain Neu5Ac or 8-keto-Neu5Ac, the overall surface charge (minus polySia) and protection of underlying galactose resides is not substantially affected, in contrast to what was observed for previous sialyltransferase inhibitors. The observation that the polysialyltransferase ST8SiaIV also had reduced efficiency with 8-keto-Neu5Ac *in vitro* is not likely to substantially affect the end goal of polySia reduction as only one addition of 8-keto-Neu5Ac is necessary to prevent further elongation.

In the past, specific reduction of α-2,8-linked Neu5Ac glycoforms required either genetic manipulation of the ST8 family sialyltransferases or enzymatic treatment of cells with the polySia hydrolase, EndoN. Unfortunately, genetic manipulation of primary cells with viral vectors or transfection reagents suffers from low efficiencies and potential toxicity issues. Similarly, the polySia hydrolase is expressed in *E. coli* and so comes with its own toxicity issues as it can be contaminated with endotoxin, complicating experiments with primary immune cells which have TLR4, including T and dendritic cells [42, 43]. Treatment with 8-keto-Neu5Ac resulted in a substantial decrease in polySia in primary human T cells in as little as 24 h, with no obvious toxicity issues. Additionally, few methods allow for the temporal control of α-2,8-linked sialyl glycoforms in primary cells such as was achieved with 8-keto-Neu5Ac, which will be useful for studying the role of polySia in biological processes at different time scales.

With 8-keto-Neu5Ac, it will now be possible to investigate the biological role of polySia in greater detail than ever before. For example, the role of polySia in immune-mediated cancer cell killing over the course of several days could be investigated. Compound **3**, like its di-fluoro-Neu5Ac analog, is likely to have good bioavailability [30], which would facilitate experiments in mice, allowing us to determine the contribution of polySia to various stages of metastasis. In addition to polySia, α2,8-linked Neu5Ac also occurs as part of the b- and c-series gangliosides, which are drug targets for several different types of cancer [44–46], suggesting further uses for the reported inhibition strategy.

## Methods and Materials

### Synthesis of 8-keto-Neu5Ac derivatives

Per-*O*-acetylated 8-keto-Neu5Ac esters were synthesized as previously described [29, 30] (See SI methods for details). For synthesis of CMP-8-keto-NeuAc, the CMP-Neu5Ac synthetase from *Neisseria meningitidis* was purified as previously described [36] (See SI methods for details).

### Cell lines

The MCF-7 cell line (ATCC HTB-22) was purchased from ATCC and was maintained in D-MEM/F-12 medium supplemented with 10% FBS and 1X penicillin-streptomycin. AtT-20 was purchased from ATCC (CCL-89) and maintained in F-12K medium supplemented with 2.5% FBS, 15% horse serum and 1X penicillin-streptomycin. Cells were grown in a humidified incubator at 37°C with 5% CO_2_.

### Toxicity assay

MCF7 cells were split into a 96-well plate at 2000 cells well and incubated overnight before addition of 2-fold serial dilutions of 8-keto-NeuAc compounds in fresh media. Cells were incubated with 8-keto-NeuAc derivatives for 24 h and proliferation measured using WST-1 (Roche Applied Science) according to manufacturer’s instructions. AtT-20 cells were split 1:2 into serially diluted 8-keto-NeuAc compounds and proliferation was measured as for MCF7 cells.

### Flow cytometry

#### Labeling cell surface ketones

MCF-7 cells were incubated with 100 μM **2 – 4** for two days. Cells were washed twice in PBS and lifted using StemPro Accutase (Gibco), then labeled with biotin-hydrazide and FITC-avidin according to Yarema et al J. Biol. Chem. 1998 273(47):31168-31179 [47].

#### Comparison of 8-keto-Neu5Ac and 3-fluoro-Neu5Ac

MCF7 cells were seeded at approximately 20 % confluency in DMEM-F12 1:1 media supplemented with 1X penicillin/streptomycin and 10 % FBS. After 1 day, the media was removed and new media containing 200 μM peracetylated 8-keto sialic acid pentyl ester, 200 μM peracetylated 3-fluoro sialic acid pentyl ester, or an equivalent volume of DMSO were added to the cells. After 3 days, cells were harvested using 1mM EDTA in PBS. Cells were washed 2 times with wash buffer (PBS with 0.5 % BSA, 0.1 mM CaCl_2_) then resuspended in 100 μL wash buffer with 10 μg/mL biotinylated MAL II (Vector Biolabs B-1265) and incubated on ice for 30 minutes. Cells were washed 2 times then resuspended in 100 μL wash buffer with 0.1 μg/mL streptavidin-PE (Biolegend 405203) and incubated on ice for 15 min. After washing cells 2 times cells were filtered before flow cytometry analysis (Cytek Aurora)

### Immunoblotting

Cells were resuspended in RIPA buffer supplemented with 1 mM PMSF and benzonase and lysed at 4°C for 30 min. Cell debris was pelleted at 18 000 ×*g* for 15 min at 4°C. The protein concentration of the supernatant was measured using a BCA assay (Pierce) and equivalent amounts of protein were loaded on SDS-PAGE gels and transferred to PVDF membrane. Membranes were blocked in 5% BSA, 1X TBS-tween, then incubated with 1:1000 α-polySia mAb 735 (BioAspect) or 1:2000 α-NCAM (R&D Systems) in blocking buffer at room temperature overnight. After washing in 1X TBS-tween, membranes were incubated with 1:1000 secondary conjugate-HRP for 1 h, then washed and developed using Pierce ECL western blotting substrate.

### Enzyme assays

ST3GalI and ST6GalI were purified as previously described [48]. ST3GalI activity was measured in reactions containing 0.5 mM T-antigen-BODIPY, 50 mM NaHEPES pH 7.4, 10 mM MnCl_2_, 1.5 mM donor sugar. ST6GalI activity was measured in reactions containing 0.5 mM LacNAc-BODIPY, 50 mM sodium phosphate pH 6.8, 10 mM MgCl_2_, 1.5 mM donor sugar. In mixed donor sugar assays, both the natural and modified sugars were present at 1.5 mM each. All reactions were incubated at 37 °C and 5 μL aliquots were stopped with an equal volume of acetonitrile at appropriate times. Product formation was quantified by HPLC using a Roc C18 column (3 μm, 3.0 × 100 mm, Restek Corporation, Bellefonte, PA) in 10 mM ammonium acetate pH 5.5, with a gradient of 25 – 45% acetonitrile over 5 CV, and fluorometric detection at excitation wavelength of 503 nm and emission wavelength of 514 nm. ST8SiaIV activity was measured in reactions containing 0.5 mM disialyl-Lac-BODIPY, 50 mM Na HEPES pH 7, 10 mM MnCl_2_, and 2 mM donor sugar. Timepoints were taken at 0 and 4 hours. The reaction was quenched with an equivalent volume of acetonitrile. BODIPY-tagged products were separated on DNAPac PA-100 Guard with 20 % acetonitrile, 0-50 % 2 M NH_4_OAc pH 8 over 20 minutes with an excitation wavelength of 503 nm and emission wavelength of 514 nm.

### T cell assays

CD3^+^ T cells were isolated from human PBMCs (Stemcell) using a negative selection kit (Stemcell) and then cultured in Immunocult media (Stemcell) supplemented with 1X penicillin-streptomycin and 25 U/mL IL-2. Cells were activated with Immunocult™ human CD3/CD28/CD2 T cell activator (Stemcell). At 6 days post-activation cells were diluted to 5 × 10^5^ cells/mL and were treated with or without 200 μM **3**. After 3 days in culture with **3** cells were diluted to 5 × 10^5^ cells/mL with fresh media and were cultured for 2 more days. Cells were harvested each day and cell pellets were frozen for future immunoblotting. Immunoblotting was performed as described above.

## Acknowledgements

We would like to acknowledge the following funding sources: GlycoNet grant to LW, NSERC grants to LW, SW, and MN, CIHR grant to WWW, NCJS and SW. We thank Dr. Lyann Sim for helpful discussions.

## Notes

### Competing Interest Statement

The authors have declared no competing interest.

## References

1. Varki, A., Essentials of glycobiology. Third edition. ed. 2017, Cold Spring Harbor, New York: Cold Spring Harbor Laboratory Press. xxix, 823 pages.

2. Varki, A., Glycan-based interactions involving vertebrate sialic-acid-recognizing proteins. Nature, 2007. 446(7139): p. 1023–1029.

3. Zarbock, A., et al., Leukocyte ligands for endothelial selectins: specialized glycoconjugates that mediate rolling and signaling under flow. Blood, 2011. 118(26): p. 6743–51.

4. Xiong, X., J.W. McCauley, and D.A. Steinhauer, Receptor binding properties of the influenza virus hemagglutinin as a determinant of host range. Curr Top Microbiol Immunol, 2014. 385: p. 63–91.

5. Stanley, P., What Have We Learned from Glycosyltransferase Knockouts in Mice? J Mol Biol, 2016. 428(16): p. 3166–3182.

6. Falconer, R.A., et al., Polysialyltransferase: a new target in metastatic cancer. Curr Cancer Drug Targets, 2012. 12(8): p. 925–39.

7. Colley, K.J., K. Kitajima, and C. Sato, Polysialic acid: biosynthesis, novel functions and applications. Crit Rev Biochem Mol Biol, 2014. 49(6): p. 498–532.

8. Yang, W.H., et al., Coordinated roles of ST3Gal-VI and ST3Gal-IV sialyltransferases in the synthesis of selectin ligands. Blood, 2012. 120(5): p. 1015–26.

9. Wang, L.B., et al., Sialyltransferase inhibition and recent advances. Biochimica Et Biophysica Acta-Proteins and Proteomics, 2016. 1864(1): p. 143–153.

10. Volkers, G., et al., Structural Basis for Binding of Fluorescent CMP-Neu5Ac Mimetics to Enzymes of the Human ST8Sia Family. ACS Chemical Biology, 2018. 13(8): p. 2320–2328.

11. Preidl, J.J., et al., Fluorescent Mimetics of CMP-Neu5Ac Are Highly Potent, Cell-Permeable Polarization Probes of Eukaryotic and Bacterial Sialyltransferases and Inhibit Cellular Sialylation (vol 53, pg 5700, 2014). Angewandte Chemie-International Edition, 2014. 53(31).

12. Miyazaki, T., et al., CMP substitutions preferentially inhibit polysialic acid synthesis. Glycobiology, 2008. 18(2): p. 187–94.

13. Macauley, M.S., et al., Systemic Blockade of Sialylation in Mice with a Global Inhibitor of Sialyltransferases. Journal of Biological Chemistry, 2014. 289(51): p. 35149–35158.

14. Rillahan, C.D., et al., Global metabolic inhibitors of sialyl- and fucosyltransferases remodel the glycome. Nat Chem Biol, 2012. 8(7): p. 661–8.

15. Wratil, P.R., et al., A Novel Approach to Decrease Sialic Acid Expression in Cells by a C-3-modified N-Acetylmannosamine. Journal of Biological Chemistry, 2014. 289(46): p. 32056–32063.

16. Curreli, S., et al., Polysialylated neuropilin-2 is expressed on the surface of human dendritic cells and modulates dendritic cell-T lymphocyte interactions. J Biol Chem, 2007. 282(42): p. 30346–56.

17. Stamatos, N.M., et al., Changes in polysialic acid expression on myeloid cells during differentiation and recruitment to sites of inflammation: role in phagocytosis. Glycobiology, 2014. 24(9): p. 864–79.

18. Shahraz, A., et al., Anti-inflammatory activity of low molecular weight polysialic acid on human macrophages. Sci Rep, 2015. 5: p. 16800.

19. Heimburg-Molinaro, J., et al., Cancer vaccines and carbohydrate epitopes. Vaccine, 2011. 29(48): p. 8802–26.

20. Gong, L., et al., Effects of the regulation of polysialyltransferase ST8SiaII on the invasiveness and metastasis of small cell lung cancer cells. Oncol Rep, 2017. 37(1): p. 131–138.

21. Tanaka, F., et al., Expression of polysialic acid and STX, a human polysialyltransferase, is correlated with tumor progression in non-small cell lung cancer. Cancer Res, 2000. 60(11): p. 3072–80.

22. Al-Saraireh, Y.M., et al., Pharmacological inhibition of polysialyltransferase ST8SiaII modulates tumour cell migration. PLoS One, 2013. 8(8): p. e73366.

23. Elkashef, S.M., et al., Polysialic acid sustains cancer cell survival and migratory capacity in a hypoxic environment. Sci Rep, 2016. 6: p. 33026.

24. Suzuki, M., et al., Polysialic acid facilitates tumor invasion by glioma cells. Glycobiology, 2005. 15(9): p. 887–94.

25. Schreiber, S.C., et al., Polysialylated NCAM represses E-cadherin-mediated cell-cell adhesion in pancreatic tumor cells. Gastroenterology, 2008. 134(5): p. 1555–66.

26. Daniel, L., et al., A nude mice model of human rhabdomyosarcoma lung metastases for evaluating the role of polysialic acids in the metastatic process. Oncogene, 2001. 20(8): p. 997–1004.

27. Scheidegger, E.P., et al., In vitro and in vivo growth of clonal sublines of human small cell lung carcinoma is modulated by polysialic acid of the neural cell adhesion molecule. Lab Invest, 1994. 70(1): p. 95–106.

28. Morley, T.J. and S.G. Withers, Chemoenzymatic Synthesis and Enzymatic Analysis of 8-Modified Cytidine Monophosphate-Sialic Acid and Sialyl Lactose Derivatives. Journal of the American Chemical Society, 2010. 132(27): p. 9430–9437.

29. Chung, K. and R.M. Waymouth, Selective Catalytic Oxidation of Unprotected Carbohydrates. Acs Catalysis, 2016. 6(7): p. 4653–4659.

30. Arns, S., et al., Assessing the oral bioavailability of difluorosialic acid prodrugs, potent viral neuraminidase inhibitors, using a snapshot PK screening assay. Bioorg Med Chem Lett, 2015. 25(12): p. 2505–9.

31. Martersteck, C.M., et al., Unique alpha 2, 8-polysialylated glycoproteins in breast cancer and leukemia cells. Glycobiology, 1996. 6(3): p. 289–301.

32. Alcaraz, G. and C. Goridis, Biosynthesis and processing of polysialylated NCAM by AtT-20 cells. Eur J Cell Biol, 1991. 55(1): p. 165–73.

33. Volkers, G., et al., Structural Basis for Binding of Fluorescent CMP-Neu5Ac Mimetics to Enzymes of the Human ST8Sia Family. ACS Chem Biol, 2018. 13(8): p. 2320–2328.

34. Mahal, L.K., et al., A small-molecule modulator of poly-alpha 2,8-sialic acid expression on cultured neurons and tumor cells. Science, 2001. 294(5541): p. 380–382.

35. Pon, R.A., N.J. Biggs, and H.J. Jennings, Polysialic acid bioengineering of neuronal cells by N-acyl sialic acid precursor treatment. Glycobiology, 2007. 17(3): p. 249–260.

36. Karwaski, M.F., W.W. Wakarchuk, and M. Gilbert, High-level expression of recombinant Neisseria CMP-sialic acid synthetase in Escherichia coli. Protein Expr Purif, 2002. 25(2): p. 237–40.

37. Collins, B.E. and J.C. Paulson, Cell surface biology mediated by low affinity multivalent protein-glycan interactions. Current Opinion in Chemical Biology, 2004. 8(6): p. 617–625.

38. Ulloa, F. and F.X. Real, Differential distribution of sialic acid in alpha2,3 and alpha2,6 linkages in the apical membrane of cultured epithelial cells and tissues. J Histochem Cytochem, 2001. 49(4): p. 501–10.

39. Xia, J., et al., Synthesis of fluorinated mucin core 2 branched oligosaccharides with the potential of novel substrates and enzyme inhibitors for glycosyltransferases and sulfotransferases. Journal of Organic Chemistry, 2006. 71(10): p. 3696–3706.

40. Hull, R.L., et al., Inhibition of glycosaminoglycan synthesis and protein glycosylation with WAS-406 and azaserine result in reduced islet amyloid formation in vitro. American Journal of Physiology-Cell Physiology, 2007. 293(5): p. C1586–C1593.

41. Buschmann, H.J., E. Dutkiewicz, and W. Knoche, The Reversible Hydration of Carbonyl-Compounds in Aqueous-Solution .2. The Kinetics of the Keto Gem-Diol Transition. Berichte Der Bunsen-Gesellschaft-Physical Chemistry Chemical Physics, 1982. 86(2): p. 129–134.

42. Reynolds, J.M., et al., Toll-like receptor 4 signaling in T cells promotes autoimmune inflammation. Proceedings of the National Academy of Sciences of the United States of America, 2012. 109(32): p. 13064–13069.

43. McAleer, J.P. and A.T. Vella, Understanding how lipopolysaccharide impacts CD4 T-cell immunity. Critical reviews in immunology, 2008. 28(4): p. 281–299.

44. Sphyris, N., et al., GD2 and GD3 synthase: novel drug targets for cancer therapy. Mol Cell Oncol, 2015. 2(3): p. e975068.

45. Sarkar, T.R., et al., GD3 synthase regulates epithelial-mesenchymal transition and metastasis in breast cancer. Oncogene, 2015. 34(23): p. 2958–67.

46. Yeh, S.C., et al., Glycolipid GD3 and GD3 synthase are key drivers for glioblastoma stem cells and tumorigenicity. Proc Natl Acad Sci U S A, 2016. 113(20): p. 5592–7.

47. Yarema, K.J., et al., Metabolic delivery of ketone groups to sialic acid residues. Application To cell surface glycoform engineering. J Biol Chem, 1998. 273(47): p. 31168–79.

48. Janesch, B., et al., Comparison of α2,6-sialyltransferases for sialylation of therapeutic proteins. Glycobiology, 2019. 29(10): p. 735–747.

