## Supplemental for "Attenuation of polysialic acid biosynthesis in cells by the small molecule inhibitor 8-keto-sialic acid"

### 1. Synthesis of titration reagents

**General Synthetic Procedures.** All reactions involving air or moisture sensitive reactants were conducted under a positive pressure of dry argon. All solvents and chemicals were reagent grade and used as supplied unless otherwise stated. For anhydrous reactions, solvents were dried according to the procedures detailed in Perrin and Armarego[2]. Removal of solvent was performed under reduced pressure, below 40 °C, using a Büchi rotary evaporator. All other chemical reagents were purchased from *Sigma-Aldrich Chemical Company*. All reactions and fractions from column chromatography were monitored by thin layer chromatography (TLC). Analytical TLC was done on glass plates (5 × 1.5 cm) precoated (0.25 mm) with silica gel (normal $SiO_2$, Merck 60 F254). Compounds were visualized by exposure to UV light and by dipping the plates in 1% $Ce(SO_4)_2 \bullet 4H_2O$ 2.5% $(NH_4)Mo_7O_{24} \bullet 4H_2O$ in 10% $H_2SO_4$ followed by heating on a hot plate. Flash chromatography was performed on silica gel (EM Science, 60Å, 230-400 mesh). The NMR spectra were either recorded on a Bruker Av-600 (600 MHZ, with Cryoprobe), Bruker AV-400 (400 MHz) or a Bruker AV-300 (300 MHz) spectrometer. Mass spectra were recorded by using a Waters/Micromass instrument (electrospray ionization) and recorded using an ion-trap.

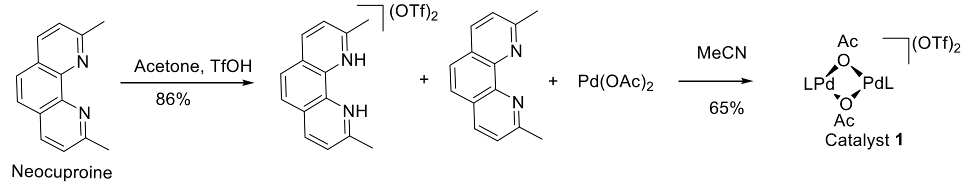


**Scheme 1**. Synthesis of Pd catalyst **1**.

**[(2,9-Dimethyl-1,10-phenanthroline)Pd(μ-OAc)]$^2$ (OTf)$^2$** (**1**): Catalyst **1** was prepared following a literature procedure with slightly modification[1]. In a 250-mL round-bottom flask was added neocuproine (3.15 g, 15.0 mmol), followed by 75 mL of acetone. Triflic acid (2.65 mL, 30.0 mmol) was then added. The resulting solution was stirred at room temperature for 50 min, and 60 mL $Et_2O$ was added. The resulting precipitates was filtered to give a white solid 5.40 g. In a 500-mL round bottle flask was added $Pd(OAc)_2$ (4.75 g, 21.2 mmol) and 50 mL MeCN. The product (5.40 g) from first step in 50 mL

MeCN and 2.20 g neocuproine (10.58 mmol) in 50 mL MeCN were added at the same time. The resulting solution was stirred at room temperature for 12 h. $Et_2O$ (300 mL) was added, and the resulting precipitates was filtered to get desired product as orange solid (6.2 g, 56%).

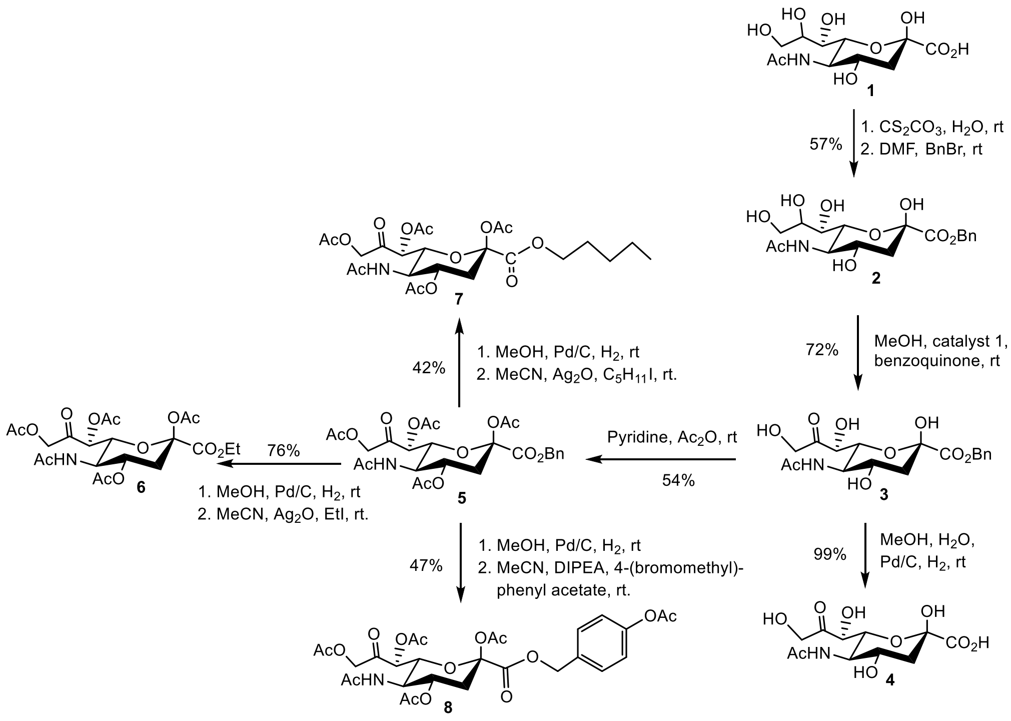

**Scheme 2**: Synthesis of 8-keto-sialic acid ester

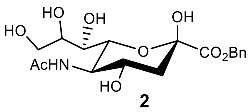

**Benzyl 5-acetamido-3,5-dideoxy-D-glycero-β-D-galacto-2-nonulopyranosonate (2)** The known compound **2** was synthesized following a literature procedure with some modification[2]. To a stirring solution of Neu5Ac (5.00 g, 16.2 mmol) in water (20 mL), 10% $Cs_2CO_3$ was added dropwise to adjust the pH value of the solution to neutral (about 30 mL). The reaction mixture was then evaporated and freeze dried to get a white powder. The obtained solid was dissolved in *N,N*-dimethylformamide (20 mL), and benzyl bromide (3.00 mL) was added dropwise. The mixture was stirred under argon overnight at room temperature and then filtered through Celite. The filtrate was evaporated to a residue, and it was dissolved in EtOAc/MeOH = 5/1. The residue was purified by flash column chromatography with EtOAc/MeOH (5/1 to 3/1) to get the product, which was then recrystallized with MeOH/$Et_2O$ to get product 1.60 g as a white solid. The mother liquor was recrystallized again with Actone/$Et_2O$ to get 2.10 g of product. Total yield is 57%. **It is important to crystalize the product! If the product was not recrystallized, following oxidation step of the C8 sialic acid will proceed very slowly!!**

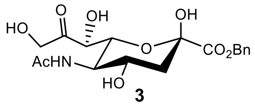


**Benzyl 5-acetamido-3,5-dideoxy-8-keto-D-glycero-β-D-galacto-2-nonulopyranoso-nate** (**3**): To a stirring solution of **2** (600 mg, 1.50 mmol) in MeOH (50 mL), Catalyst **1** (77 mg, 0.15 mmol) was added, followed by 1,4-benzoquinone (322 mg, 3.00 mmol). The solution was stirred at 25 °C for 4 h, EtOAc (20 mL) was added. The residue was concentrated down to about 3 mL, and silica gel was added and evaporated to dryness. The product was purified by flash column chromatography (EtOAc/EtOH=9/1) to afford **3** as a brownish solid (430 mg, 72%). **$^1$H NMR** (400 MHz, $D_2O$) $\delta$ 7.45 (m, 5 H, Ar-H), 4.61 (d, $J$ = 19.2 Hz, 1H, H-9), 4.43 (d, 1H, H-9), 4.39 (d, $J$ = 1.6 Hz, 1H, H-7), 4.22 (dd, $J$ = 10.4, 1.6 Hz, 1H, H-6), 4.6 (m, 1H, H-4), 3.95 (t, $J$ = 10.4 Hz, 1H, H-5), 2.29 (dd, $J$ = 12.8, 4.8 Hz, 1H, H-3), 2.05 (s, 3H), 1.91 (dd, $J$ = 11.6 Hz, 1H, H-3). **$^{13}$C NMR** (100 MHz, $D_2O$) $\delta$ 213.1 (C8), 175.0, 170.2, 135.1, 129.1 (3C), 128.5 (2C), 95.6, 74.2, 73.4, 68.5, 66.5, 51.9, 38.8, 22.3. **HRMS (ESI) m/z**: Calcd. for $C_{18}H_{23}NO_9$ ([M + H]$^+$): 398.1451; found: 398.1450.

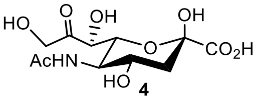


**5-acetamido-3,5-dideoxy-8-keto-D-glycero-β-D-galacto-2-nonulopyranosonate** (**4**): To a stirred solution of **3** (145 mg, 0.36 mmol) in 20 mL of MeOH/$H_2O$ = 1/1 was added 28 mg Pd/C. The solution was stirred under $H_2$ for 1 h. The solution was filtered through celite and the filtrate was concentrated to about 5 mL. The residue was loaded to a 2 g C18 column, and was eluted with water. The fraction containing the product was freeze dried to get product (111 mg, 99%) as a white solid. **$^{1}$H NMR** (400 MHz, $D_2O$) $\delta$ 4.59 (d, $J$ = 19.6 Hz, 1H, H-9), 4.48 (d, $J$ = 19.6 Hz, 1H, H-9), 4.40 (d, $J$ = 1.7 Hz, 1H, H-7), 4.18 (dd, $J$ = 10.3, 1.6 Hz, 1H, H-6), 4.05 – 4.02 (m, 1H, H-4), 3.93 (t, $J$ = 10.2 Hz, 1H, H-5), 2.26 (dd, $J$ = 12.8, 4.5 Hz, 1H, H-3), 2.02 (s, 3H), 1.85 (t, $J$ = 12.3 Hz, 1H, H-3). **$^{13}$C NMR** (100 MHz, $D_2O$) $\delta$ 213.0 (C8), 175.0, 173.0, 97.8(C2), 74.2(C7), 73.3(C6), 66.6 (C4), 66.4 (C9), 51.9(C5), 39.0(C3), 22.3. **HRMS (ESI) m/z**: Calcd. for $C_{11}H_{16}NO_9$ ([M - H]$^-$): 306.0825; found: 306.0826.

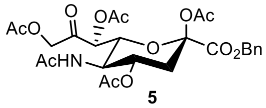


**5**

**Benzyl 5-acetamido-2,4,7,9-tetra-*O*-acetyl-3,5-dideoxy-8-keto-D-glycero-β-D-galacto-2-nonulopyranosonate** (**5**): To a stirred solution of **3** (100 mg, 0.63 mmol) in 10 mL $Ac_2O$ was added 1.0 mL pyridine. The resulting solution was stirred for 12 h at room temperature. The reaction was cooled down to 0 °C, and the reaction was quenched with the addition of MeOH (50 mL). The solvent was removed under vacuum and 50 mL of 1M HCl was added. The solution was extracted with EtOAc (50 mL x 3). The combined organic layers were washed with brine (20 mL) and dried over $Na_2SO_4$. The solvent was removed under vacuum and the residue was purified using flash column chromatography (3/1 EtOAc/ petroleum ether) to give **5** as a slightly yellow solid (158 mg, 54%). **$^{1}$H NMR** (300 MHz, $CDCl_3$) $\delta$ 7.35 (dd, $J$ = 4.8, 2.4 Hz, 5H, Ph), 5.68 (d, $J$ = 9.0 Hz, 1H, N<u>H</u>), 5.27 (d, $J$ = 2.1 Hz, 1H, H-7), 5.26 – 5.20 (m, 1H, H-4), 5.23 (d, $J$ = 12.2 Hz, 1H, C<u>H</u>$_2$Ph), 5.14 (d, $J$ = 12.3 Hz, 1H, C<u>H</u>$_2$Ph), 4.87 (d, $J$ = 17.5 Hz, 1H, H-9), 4.65 (d, $J$ = 17.5 Hz, 1H, H-9), 4.35 – 4.14 (m, 2H, H-5, H-6), 2.52 (dd, $J$ = 13.4, 5.0 Hz, 1H, H-3), 2.25 (s, 3H), 2.14 (s, 3H), 2.09 (s, 3H), 2.10 – 2.05 (m, 1H, H-3), 2.03 (s, 3H), 1.89 (s, 3H). **$^{13}$C NMR** (75 MHz, $CDCl_3$) $\delta$ 200.8, 171.2, 170.7, 170.6, 170.4, 168.2, 165.8, 135.0, 128.7, 128.7, 96.8, 75.1, 74.4, 68.0, 67.9, 67.0, 49.0, 36.3, 23.1, 21.0, 20.8, 20.7, 20.6, 20.5. **HRMS (ESI) m/z**: Calcd. for $C_{26}H_{31}NO_{13}$ ([M + Na]$^+$): 588.1693; found: 588.1671

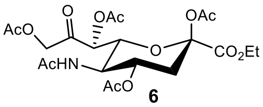

6

**Ethyl 5-acetamido-2,4,7,9-tetra-*O*-acetyl-3,5-dideoxy-8-keto-D-glycero-β-D-galacto-2-nonulopyranosonate** (**6**): To a stirred solution of **5** (195 mg, 0.35 mmol) in 12 mL of MeOH was added 15 mg of Pd/C. The solution was stirred under $H_2$ for 1 h at room temperature. The solution was filtered through a short pad of celite and the filtrate was concentrated down to a residue, which was dissolved in dry MeCN (25 mL). To the solution $Ag_2O$ (400 mg) and EtOH (0.25 mL) were added, and the suspension was stirred overnight at room temperature. The solution was filtered through a short pad of celite. The filtrate was removed under vacuum and the residue was purified using flash column chromatography (EtOAc/petroleum ether = 4/1) to give **6** as a white solid (133 mg, 76%). **$^1$H NMR** (400 MHz, $CDCl_3$) $\delta$ 5.55 (d, $J$ = 9.4 Hz, 1H, N<u>H</u>), 5.27 (d, $J$ = 2.0 Hz, 1H, H-7), 5.26 – 5.19 (m, 1H, H-4), 4.90 (d, $J$ = 17.6 Hz, 1H, H-9), 4.74 (d, $J$ = 17.5 Hz, 1H, H-9), 4.36 – 4.06 (m, 4H, H-5, H-6, OC<u>H</u>$_2$CH$_3$), 2.53 (dd, $J$ = 13.4, 5.0 Hz, 1H, H-3), 2.28 (s, 3H), 2.14 (s, 2H), 2.11 (s, 3H), 2.12 – 2.05 (m, 1H, H-5, H-3), 2.05 (s, 4H), 1.91 (s, 3H), 1.27 (t, $J$ = 7.1 Hz, 3H). **$^{13}$C NMR** (75 MHz, $CDCl_3$) $\delta$ 200.7, 171.3, 170.7, 170.6, 170.3, 168.0, 165.9, 96.9, 75.2, 74.5, 68.0, 67.0, 62.5, 49.2, 36.3, 23.3, 21.0, 20.9, 20.7, 20.6, 14.0. **HRMS (ESI) m/z**: Calcd. for $C_{21}H_{29}NO_{13}$ ([M + Na]$^+$): 526.1542; found: 526.1537.

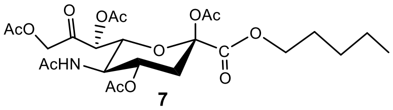

7

**Pentyl 5-acetamido-2,4,7,9-tetra-*O*-acetyl-3,5-dideoxy-8-keto-D-glycero-β-D-galacto-2-nonulopyranosonate** (**7**): To a stirred solution of **5** (175 mg, 0.37 mmol) in 12 mL of MeOH was added 14 mg of Pd/C. The solution was stirred under $H_2$ for 1 h at room temperature. The solution was filtered through a short pad of celite and the filtrate was concentrated down to a residue, and dissolved in dry MeCN (22 mL). To the solution $Ag_2O$ (360 mg) and iodopentane (0.2 mL, 1.57 mmol, 5 equiv) was added, and the suspension was stirred at room temperature overnight, then filtered through a short pad of celite. The filtrate was removed under vacuum and the residue was purified using flash column chromatography (EtOAc/petroleum ether = 3/1) to give **7** as a slightly yellow solid (71mg, 42%). **$^1$H NMR** (400 MHz, $CDCl_3$) *δ* 5.54 (d, *J* = 9.4 Hz, 1H, NH), 5.28 (d, *J* = 2.0 Hz, 1H, H-7), 5.27 – 5.22 (m, 1H, H-4), 4.92 (d, *J* = 17.5 Hz, 1H, H-9), 4.72 (d, *J* = 17.5 Hz, 1H, H-9), 4.34 – 4.11 (m, 4H, H-5, H-6, pentyl ), 2.53 (dd, *J* = 13.4, 5.0 Hz, 1H, H-3), 2.28 (s, 3H), 2.15 (s, 3H), 2.12 (s, 3H), 2.06 (s, 3H), 2.05 – 2.01 (m, 1H, H-3), 1.91 (s, 3H), 1.74 – 1.58 (m, 2H), 1.42 – 1.28 (m, 4H), 0.95 – 0.87 (m, 3H). **$^{13}$C NMR** (101 MHz, $CDCl_3$) *δ* 200.6 (C-9), 171.3, 170.7, 170.5, 170.2, 168.0, 166.0, 96.9 (C-2), 75.1, 74.5, 68.0, 66.9, 66.6, 49.3, 36.4 (C-3), 28.0, 23.2, 22.3, 21.0, 20.8, 20.7, 20.6, 14.1. **HRMS (ESI) m/z**: Calcd. for $C_{24}H_{35}NO_{13}$ ([M + Na]$^+$): 568.2006; found: 568.2000.

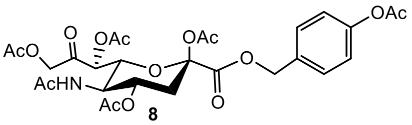

8

**4-Acetoxybenzyl 5-acetamido-2,4,7,9-tetra-*O*-acetyl-3,5-dideoxy-8-keto-D-glycero-β-D-galacto-2-nonulopyranosonate** (**8**): To a stirred solution of **5** (65.0 mg, 0.114 mmol) in 5 mL of MeOH was added 2 mg of Pd/C. The solution was stirred under $H_2$ for 1 h at room temperature. The solution was filtered through a short pad of celite and the filtrate was concentrated down to a residue. The reside was dissolved in dry MeCN (5 mL). DIPEA (0.04 mL, 0.230 mmol) and 4-(bromomethyl)phenyl acetate (79.0 mg, 0.345 mmol) were added. The solution was stirred for 12 h at room temperature. The reaction was quenched with one drop of acetic acid and the solvent was removed under vacuum and the residue was purified using flash column chromatography (EtOAc) to give **7** as a slightly yellow solid (34mg, 47%). **$^1$H NMR** (400 MHz, $CDCl_3$) *δ* 7.40 – 7.31 (m, 2H), 7.13 – 7.03 (m, 2H), 5.56 (d, *J* = 9.2 Hz, 1H, N<u>H</u>), 5.26 (d, *J* = 2.1 Hz, 1H, H-7), 5.25 – 5.20 (m, 1H, H-4), 5.22 (d, *J* = 12.3 Hz, 1H, C<u>H</u>$_2$Ph), 5.13 (d, *J* = 12.3 Hz, 1H, C<u>H</u>$_2$Ph), 4.86 (d, *J* = 17.5 Hz, 1H), 4.66 (d, *J* = 17.5 Hz, 1H), 4.25 (q, *J* = 10.0 Hz, 1H, H-5), 4.17 (dd, *J* = 10.6, 2.2 Hz, 1H, H-6), 2.51 (dd, *J* = 13.4, 5.0 Hz, 1H, H-3), 2.30 (s, 3H), 2.25 (s, 3H), 2.14 (s, 3H), 2.10 (s, 3H), 2.10 – 2.05 (m, 1H, H-3), 2.04 (s, 3H), 1.90 (s, 3H). **$^{13}$C NMR** (75 MHz, $CDCl_3$) *δ* 200.7, 170.7, 170.4, 169.4, 168.2, 165.8, 150.9, 132.6,

129.6, 121.9, 96.8, 77.6, 77.2, 76.7, 75.1, 74.4, 67.9, 67.3, 67.0, 48.9, 36.3, 29.8, 23.1, 21.2, 21.2, 21.0, 20.8, 20.6, 20.5. **HRMS (ESI) m/z**: Calcd. for $C_{28}H_{33}NO_{15}$ ($[M + Na]^+$): 646.1748; found: 646.1765.

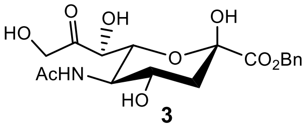
HO
O
OH
OH
O
CO2Bn
AcHN
HO

**3**

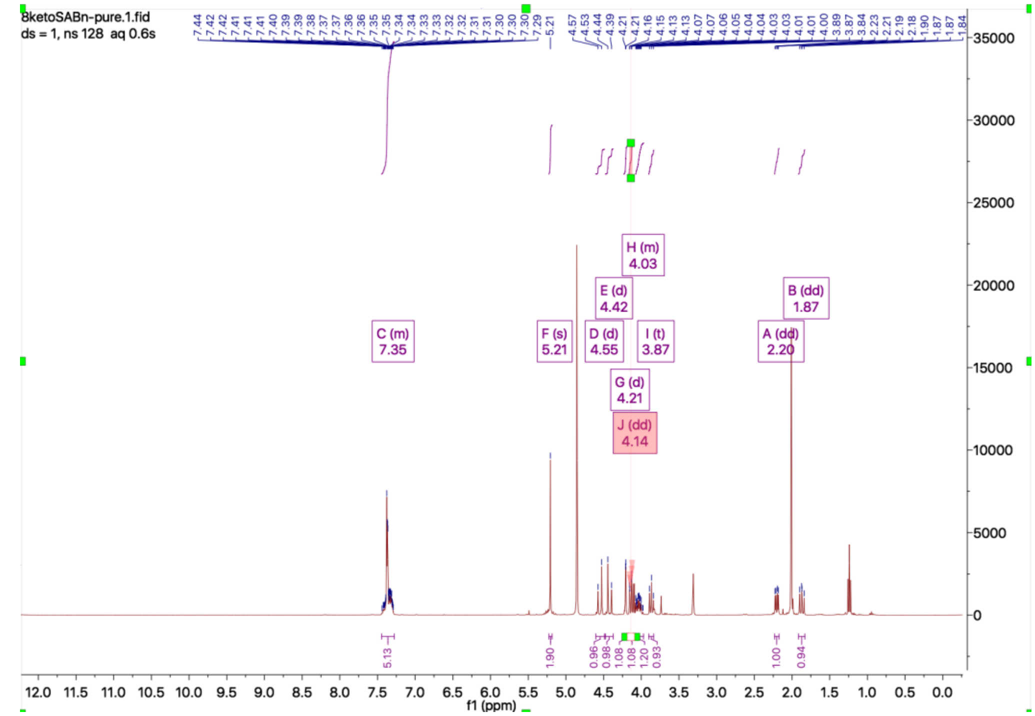
8ketoSABn-pure.1.fid
ds = 1, ns 128 aq 0.6s
H (m)
4.03
E (d)
4.42
B (dd)
1.87
C (m)
7.35
F (s)
5.21
D (d)
4.55
I (t)
3.87
A (dd)
2.20
G (d)
4.21
J (dd)
4.14
f1 (ppm)

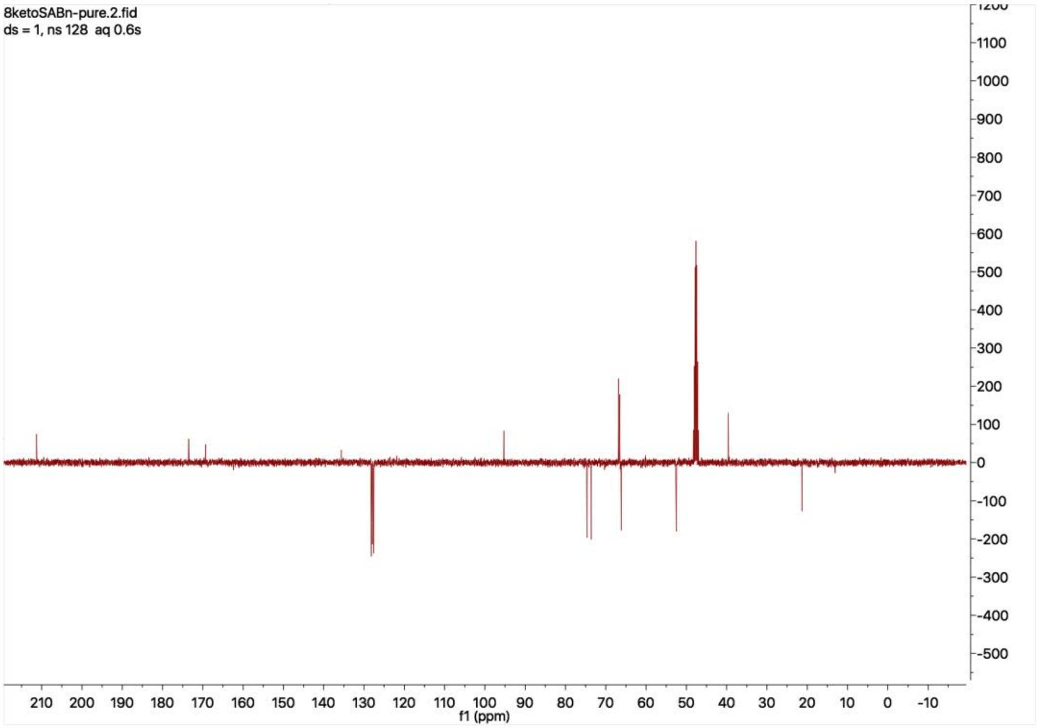
8ketoSABn-pure.2.fid
ds = 1, ns 128 aq 0.6s
f1 (ppm)

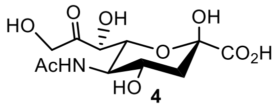


**4**

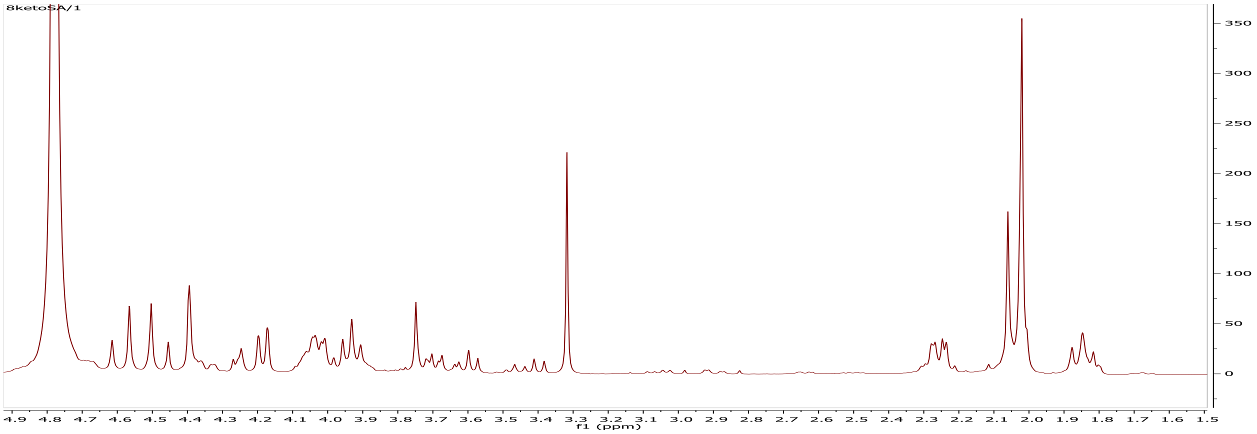


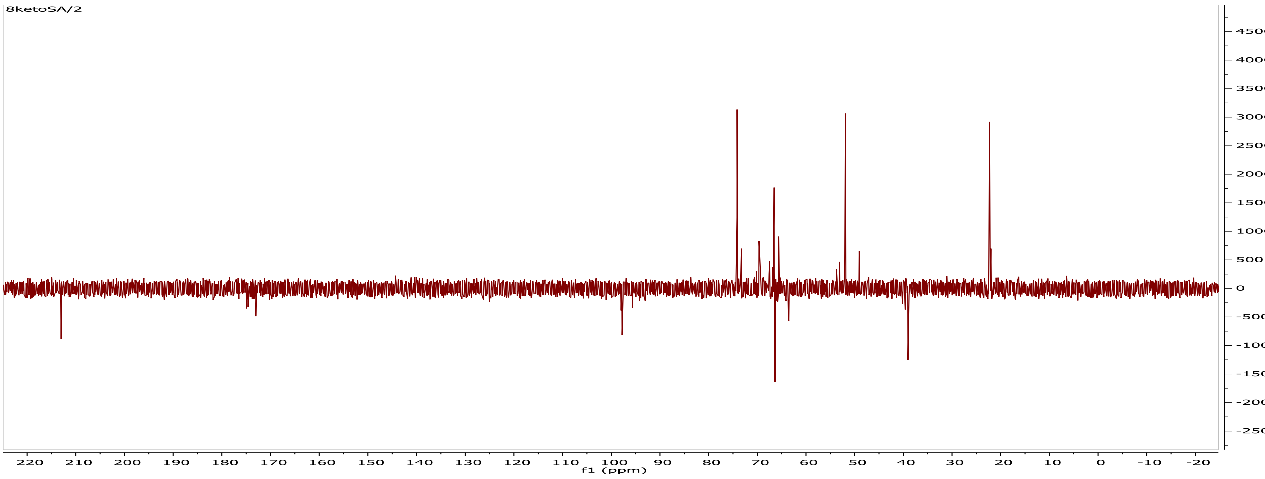

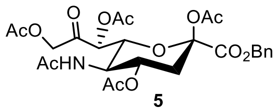
AcO
O
OAc
OAc
AcHN
O
$CO_2Bn$
AcO

**5**

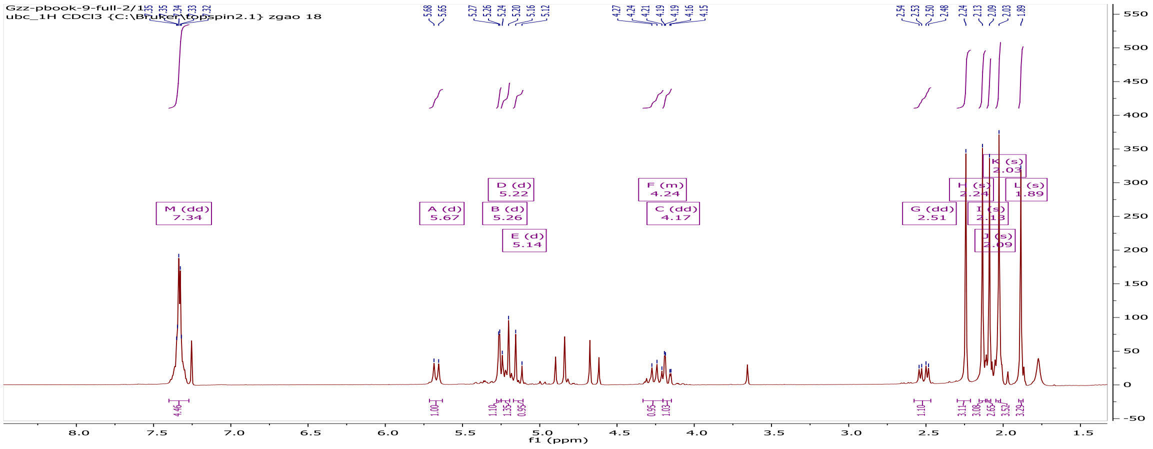
Gzz-pbook-9-full-2/1
ubc_1H CDCl3 {C:\Bruker\topspin2.1} zgao 18
M (dd) 7.34
A (d) 5.67
B (d) 5.26
D (d) 5.22
E (d) 5.14
F (m) 4.24
C (dd) 4.17
G (dd) 2.51
L (s) 1.89
f1 (ppm)

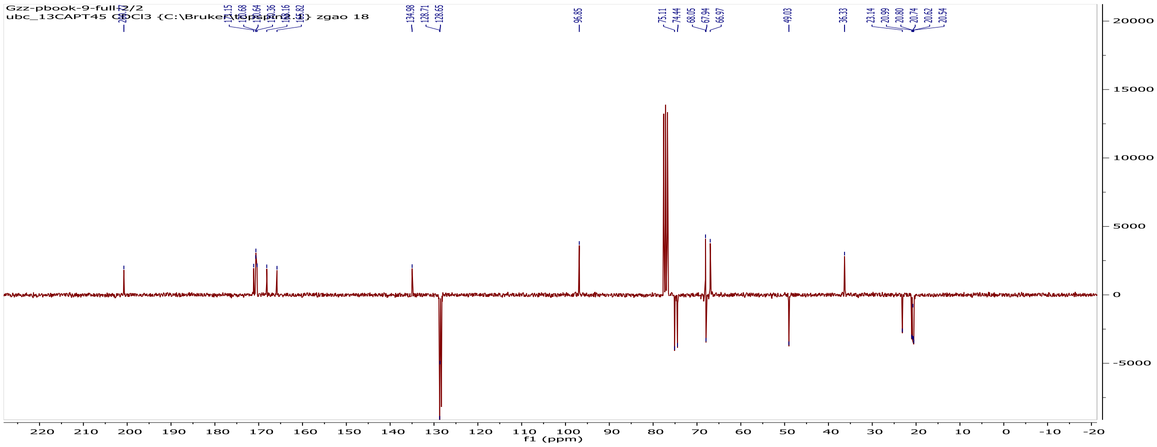
Gzz-pbook-9-full-2/2
ubc_13CAPT45 CDCl3 {C:\Bruker\topspin2.1} zgao 18
f1 (ppm)

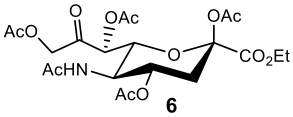
AcO
O
OAc
OAc
CO2Et
AcHN
AcO

**6**

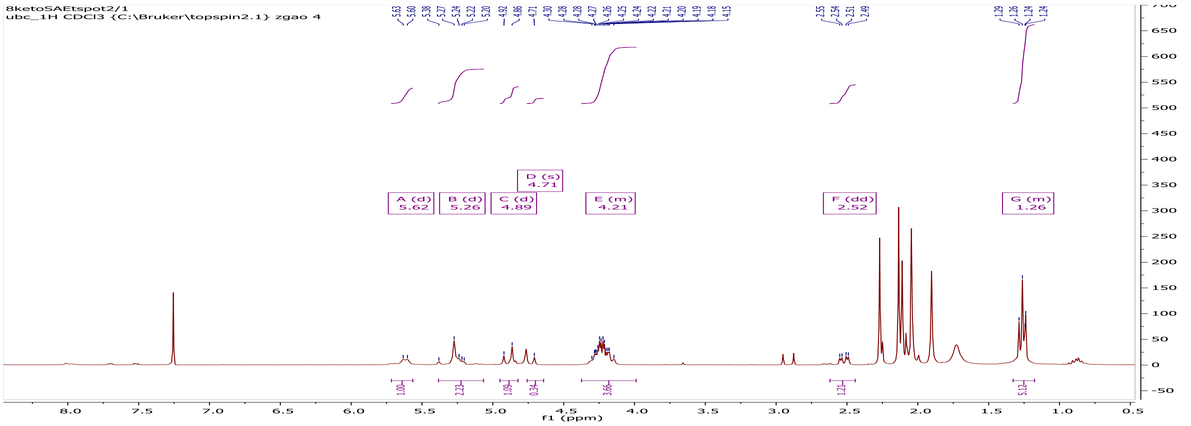
8ketoSAEtspot2/1
ubc_1H CDCl3 {C:\Bruker\topspin2.1} zgao 4
A (d) 5.62
B (d) 5.26
C (d) 4.89
D (s) 4.71
E (m) 4.21
F (dd) 2.52
G (m) 1.26
f1 (ppm)

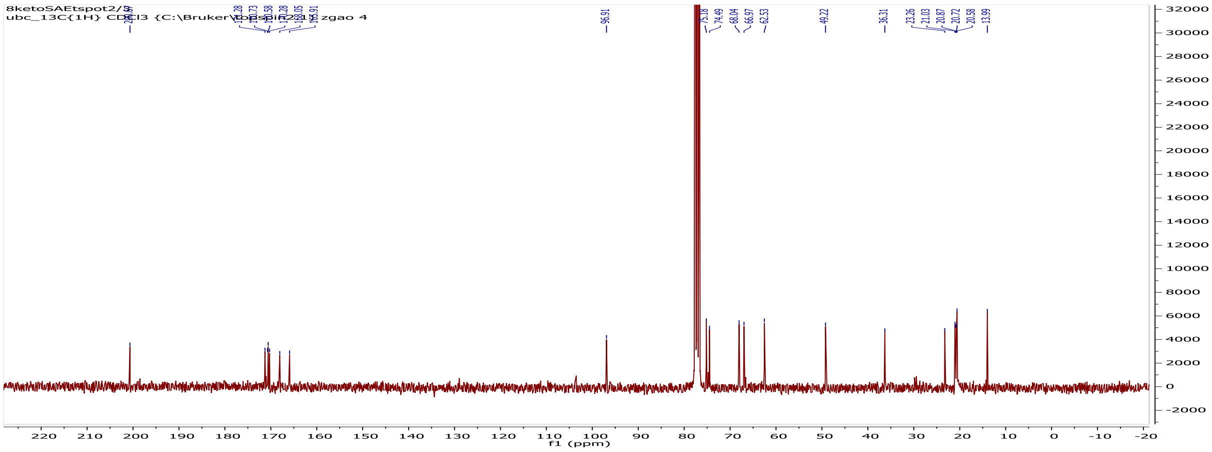
f1 (ppm)

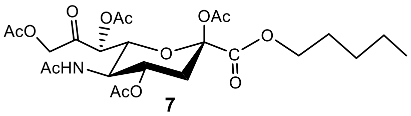
AcO
O
OAc
OAc
O
O
C
O
AcHN
AcO

**7**

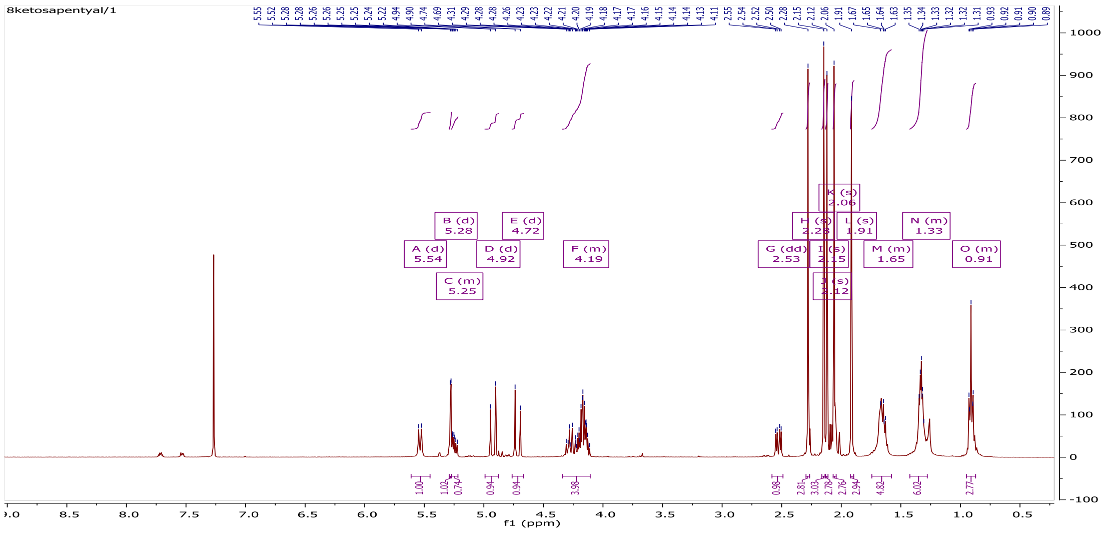
8ketosapentyal/1
A (d)
5.54
B (d)
5.28
C (m)
5.25
D (d)
4.92
E (d)
4.72
F (m)
4.19
G (dd)
2.53
M (m)
1.65
N (m)
1.33
O (m)
0.91
f1 (ppm)

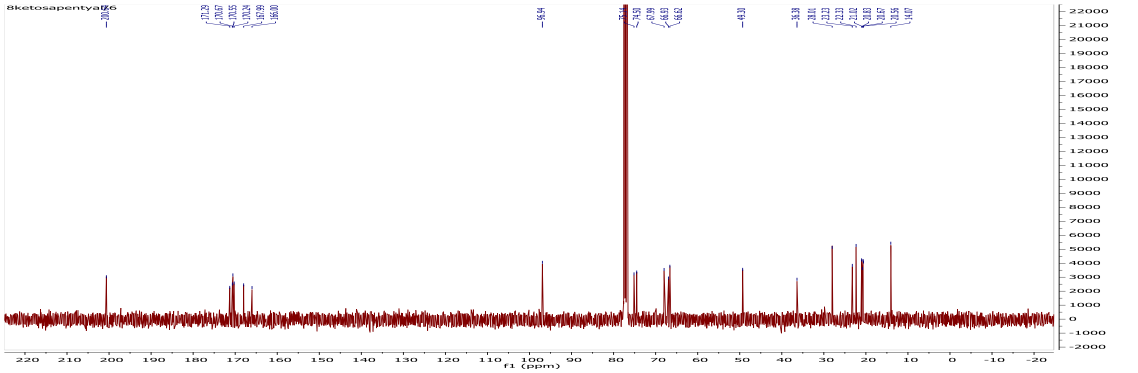
f1 (ppm)

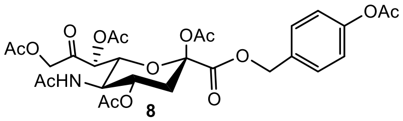

AcO
O
OAc
OAc
OAc
O
O
AcHN
AcO
O


**8**

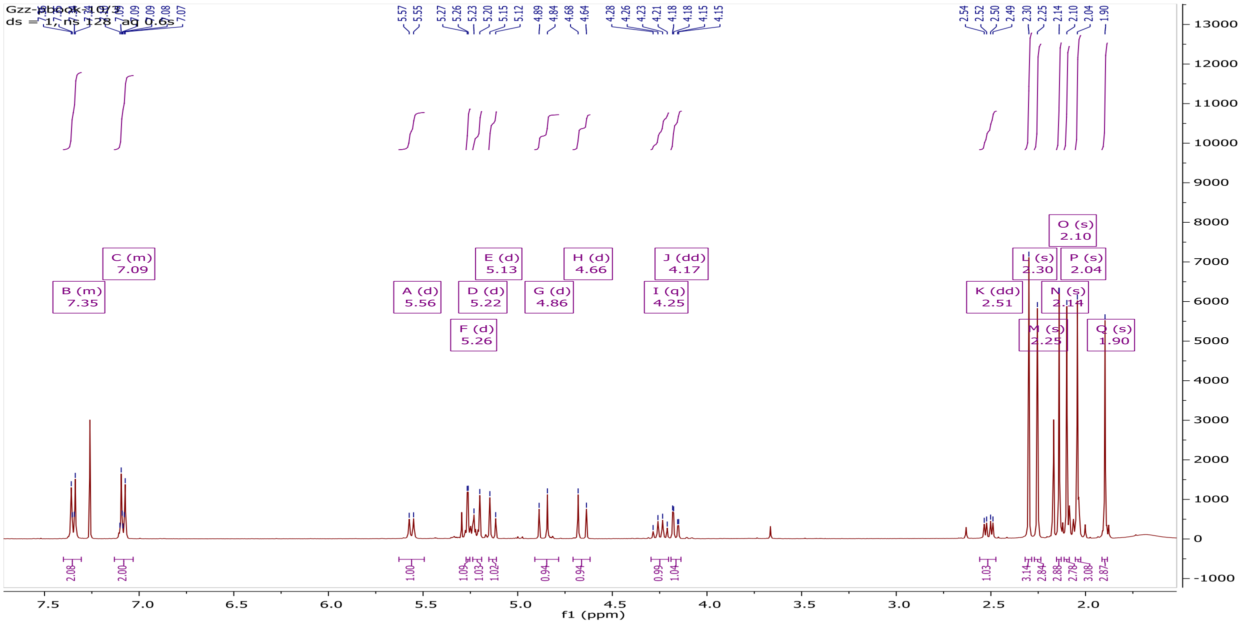

B (m)
7.35
C (m)
7.09
A (d)
5.56
D (d)
5.22
E (d)
5.13
F (d)
5.26
G (d)
4.86
H (d)
4.66
I (q)
4.25
J (dd)
4.17
K (dd)
2.51
L (s)
2.30
M (s)
2.25
N (s)
2.14
O (s)
2.10
P (s)
2.04
Q (s)
1.90
f1 (ppm)


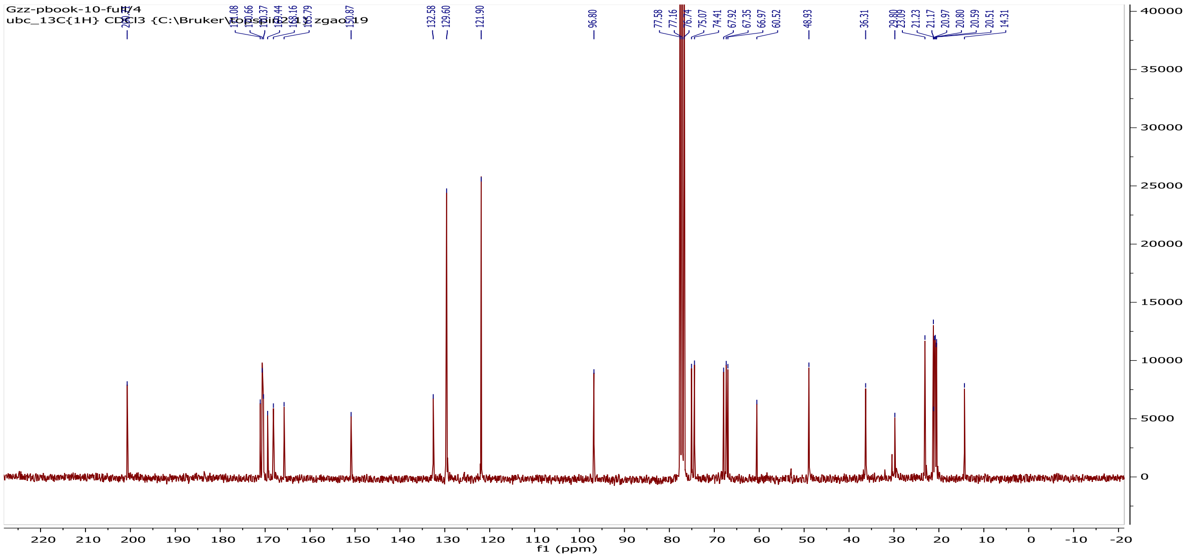

132.58
129.60
121.90
96.80
77.58
77.16
76.74
75.07
74.41
67.92
67.35
66.97
60.52
48.93
36.31
21.23
21.17
20.97
20.80
20.59
20.51
14.31
f1 (ppm)

#### 2. Additional methods

**Synthesis of CMP-8-keto-Neu5Ac**

To make CMP-8-keto-Neu5Ac, 100 mM of CTP and 8-keto-NeuAc were mixed with 200 mM TrisHCl pH 8.8, 40 mM $MgCl_2$, and 0.4 mM DTT. CMP-NeuAc synthetase was added at a final concentration of 1.5 mg/mL and the reaction was incubated at 37 °C for 2 h. Reaction progress was monitored by TLC using silica backed plates and 5:3:2 nBuOH:AcOH:$H_2O$ as the mobile phase. Once the reaction was complete, the precipitate was removed by centrifugation and the supernatant was diluted 1:30 in $H_2O$ and applied to a 5 mL HiTrap Q column. Sample was eluted with a gradient of 0 – 100% 0.5 M $NH_4HCO_3$ over 20 CV. TLC of fractions indicated that the first peak contained the desired CMP-8-keto-Neu5Ac.

#### 3. Supplemental figures

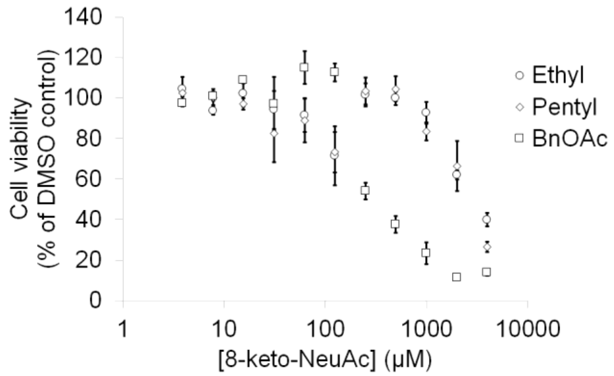


**Figure S1.** Toxicity of protected 8-keto-Neu5Ac esters in MCF7 cells, as measured using a WST-1 assay after 24 h incubation.

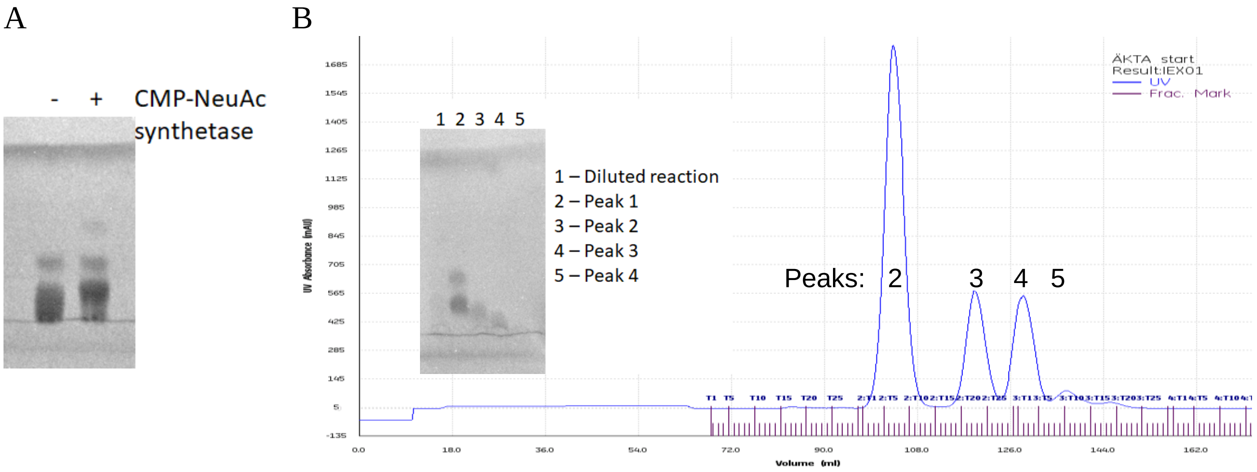


**Figure S2.** Synthesis of CMP-8-keto-Neu5Ac. A) TLC of CMP-8keto-Neu5Ac synthesis reaction. Solvent is 5:3:2 nBuOH:AcOH:$H_2O$, plastic back plate was visualized using transUV. B) Purification of CMP-8-keto-Neu5Ac. Synthesis reaction was diluted in $H_2O$ and applied to a 5 mL HiTrap Q column. Sample was eluted with a gradient of 0 – 100% 0.5 M $NH_4HCO_3$ over 20 CV. TLC of fractions indicated that Peak 2 contains CMP-8-keto-Neu5Ac.

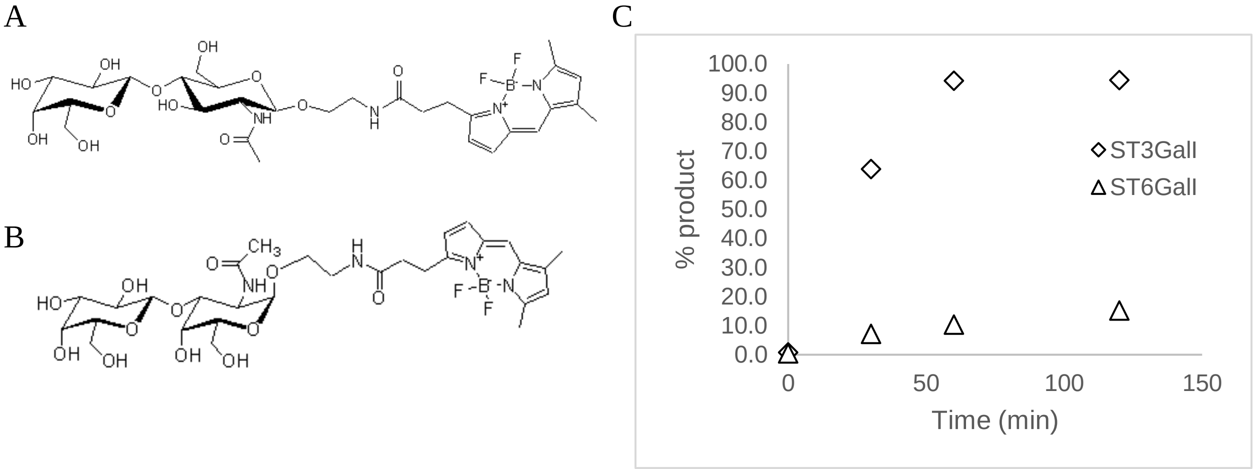


**Figure S3.** Activity of sialyltransferases with CMP-8keto-Neu5Ac. Structures of A) BODIPY-LacNAc and B) BODIPY-TAg. C) Assays are identical to those from Figure 4A.B but have increased concentration of enzyme and were performed at 37 °C. With ST3GalI, it was possible to push the reaction to completion, which meant we could then generate the data necessary for Michaelis-Menten kinetics (Figure 4C). With ST6GalI, it was not possible to push the reaction past 10% product formation.

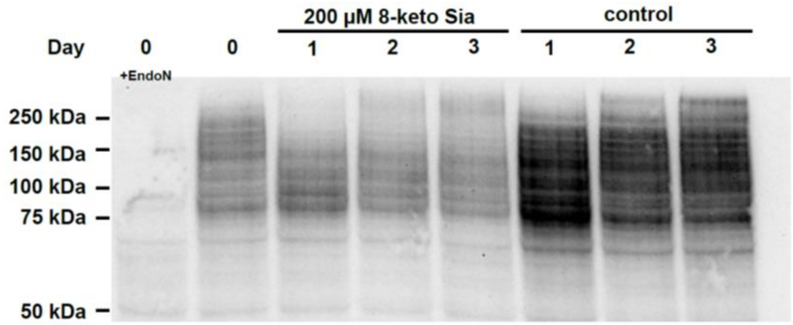


**Figure S4.** Immunoblot of polySia in primary CD3+ T cells after feeding 200 μM **3** at Day 0. Equivalent amount of protein was added into each well.

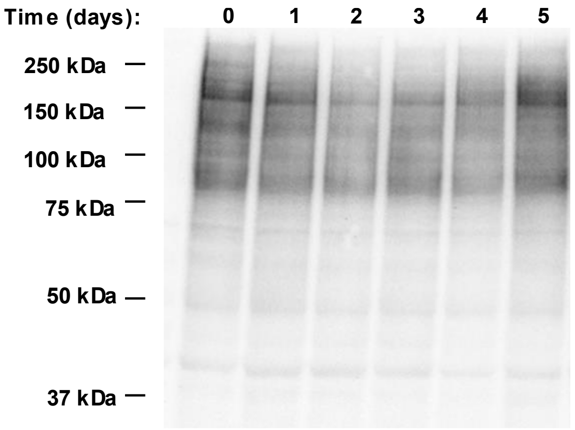


**Figure S5.** Immunoblot of polySia in MCF7 cells after feeding 100 μM **3** at day 0. Equivalent amount of protein was added into each well and 8-keto-Neu5Ac was removed from the media after 3 days.

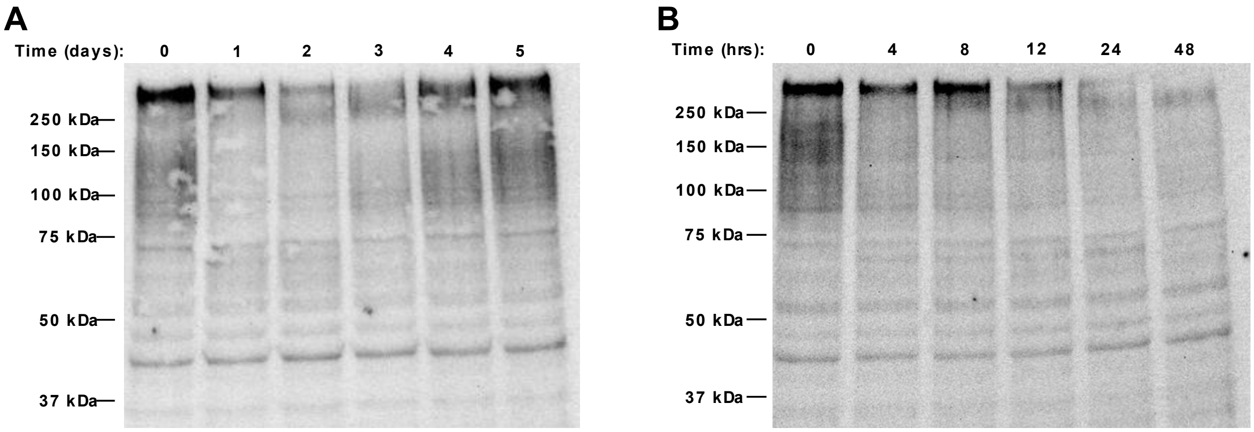


**Figure S6**. Immunoblot of polySia in Jurkat cells after feeding 100 μM **3** at day 0. Equivalent amount of protein was added into each well and 8-keto-Neu5Ac was removed from the media after 3 days.

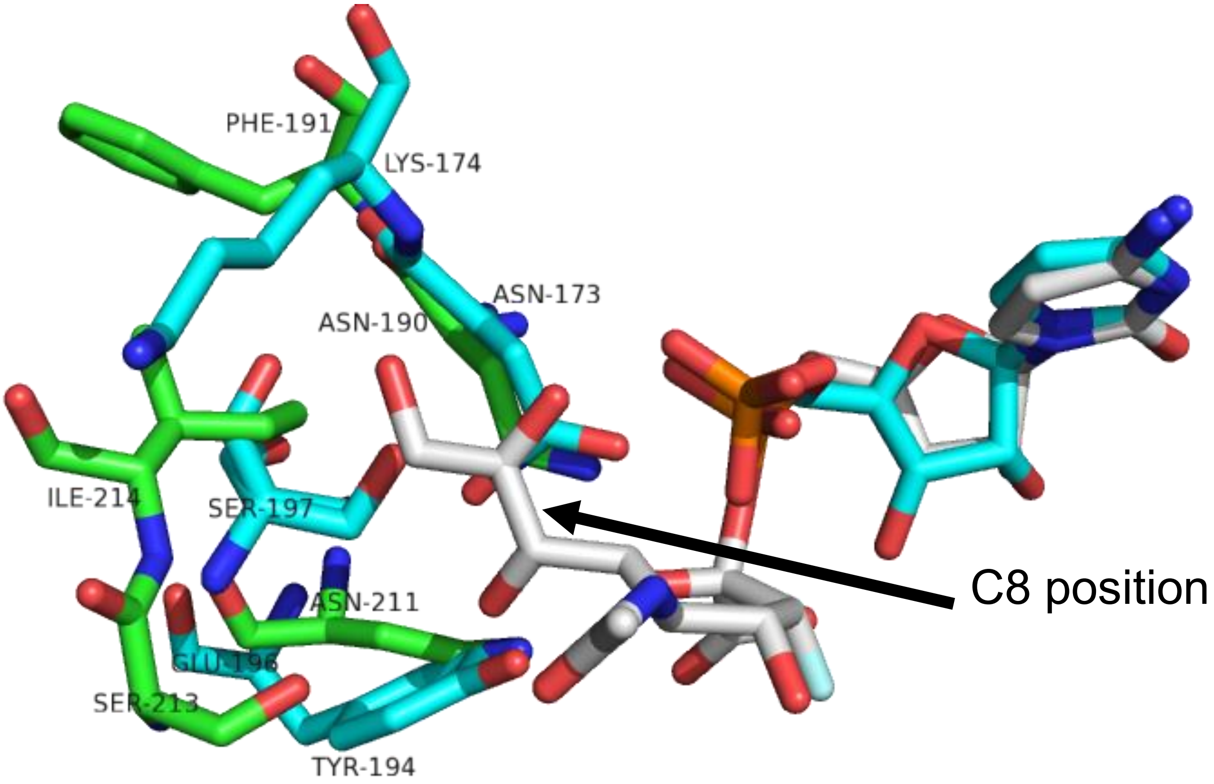


**Figure S7**. Overlay of ST8SiaIII (green) containing CMP-3-fluoro-Neu5Ac (grey) from PDB 5BO9 and ST3GalI (cyan) with CMP (cyan) from PDB 2WNB. Structures were aligned based on the CMP moiety and the conserved Asn-190/173 residue.
